# SARS-CoV-2 transcriptome analysis and molecular cataloguing of immunodominant epitopes for multi-epitope based vaccine design

**DOI:** 10.1101/2020.05.14.097170

**Authors:** Sandeep Kumar Kushwaha, Veerbhan Kesarwani, Samraggi Choudhury, Sonu Gandhi, Shailesh Sharma

## Abstract

SARS-CoV-2 is a single-stranded RNA virus that has caused more than 0.29 million deaths worldwide as of May 2020, and influence of COVID-19 pandemic is increasing continuously in the absence of approved vaccine and drug. Moreover, very limited information is available about SARS-CoV-2 expressed regions and immune responses. In this paper an effort has been made, to facilitate vaccine development by proposing multiple epitopes as potential vaccine candidates by utilising SARS-CoV-2 transcriptome data. Here, publicly available RNA-seq data of SARS-CoV-2 infection in NHBE and A549 human cell lines were used to construct SARS-CoV-2 transcriptome to understand disease pathogenesis and immune responses. In the first step, epitope prediction, MHC class I and II gene identification for epitopes, population coverage, antigenicity, immunogenicity, conservation and crossreactivity analysis with host antigens were performed by using SARS-CoV-2 transcriptome, and in the second step, structural compatibility of identified T-and B-cell epitopes were evaluated with MHC molecules and B-cell receptors through molecular docking studies. Quantification of MHC gene expression was also performed that indicated high variation in allele types and expression level of MHC genes with respect to cell lines. In A549 cell line, HLA-A*30:01:01:01 and HLA-B*44:03:01:01 were highly expressed, whereas 92 variants of HLA-A*24 genes such as HLA-A*24:02:01:01, HLA-A*24:286, HLA-A*24:479Q, HLA-A*24:02:134 and HLA-A*24:02:116 were highly expressed in NHBE cell lines. Prevalence of HLA-A*24 alleles was suggested as risk factors for H1N1 infection, and associated with type-1 diabetes. HLA-C*03:03, linked with male infertility factors was also highly expressed in SARS-CoV-2 infected NHBE cell lines. Finally, three potential T-cell and five B-cell epitopes were selected for molecular docking studies with twenty-two MHC molecules and two B-cell receptors respectively. The results of *in silico* analysis indicated that proposed epitopes have high potential to recognize immune response of SARS-CoV-2 infection. This study will facilitate *in vitro* and *in vivo* vaccine related research studies.

## 1. Introduction

Corona viruses are a group of related viruses that are responsible for causing diseases ranging from the common cold to severe diseases like Middle East respiratory syndrome (MERS), SARS-CoV-2 in mammals and birds. SARS-CoV-2 is a positive sense single-stranded RNA virus belonging to the family *Coronaviridae* and subgenus *Sarbecovirus*. It is responsible for the widespread global pandemic causing an upper respiratory tract infection of humans [1]. SARS-CoV-2 virion ranges from approximately 50-200 nm in diameter [2]. SARS-CoV-2 is made up of four structural proteins known as the S (spike), E (envelope), M (membrane) and N (nucleocapsid) proteins. The nucleocapsid protein contains the viral RNA and the spike, membrane, envelope make up the viral envelope. The spike protein is responsible for the viral attachment with angiotensin-converting enzyme 2 (ACE2) receptors and facilitates entry into the host cells [3]. The ACE2 receptors are present in the goblet (secretory) cells of ciliated cells in the nose, back of the throat, lungs, gut, heart muscles and kidney which facilitates the hand to mouth transmission route. The viral RNA is released in the nasal cells when the transmembrane serine protease 2 (TMPRSS2) splits the spike proteins, and enters inside the cell, the viral genetic material replicates into millions. Seroconversion of SARS-CoV-2 took place within four days of infection and was found in most patients by day 14 and persistent specific IgG and antibody production was reported even after 2 years of infection [4]. Whereas limited serological details of SARS-CoV-2 are available at the moment, it is reported that a patient showed the presence of IgM after 9 days of infection, and later production of IgG after 2 weeks [5]. In an *in vitro* plaque testing with patient sera, it was confirmed that it is able to neutralize SARS-CoV highlighting the successful mounting of humoral response [6]. The current evidences have shown that Th1 immune response can be successful for controlling SARS-CoV and may work for SARS-CoV-2 as well, since the epitopes overlap for both, the T-epitopes can be identified and will be valuable for designing the cross-reactive vaccines.

Epitopes are the antigenic regions of an antigen, causing an immune response which is identified by antibodies generated from T-and B-cells. T-cell epitopes present on the cell surface binds to the major histocompatibility complex (MHC) molecules. The MHC I molecules presents peptides of 8-11 amino acids in length, which are CD8^+^ T-cell epitopes. In contrast, MHC II molecules present longer peptides of 13-17 amino acids in length, which are CD4^+^ T-cell epitopes. The epitope-based vaccine development offers prospective advantages over the whole protein approach because the immune response against highly reserved epitopes over a widespread population can be used for the treatment of highly variable pathogens [7, 8]. Various successful studies were reported for the epitope-based vaccine design against West Nile virus [9], dengue virus [10], chikungunya virus [11], shigellosis [12] etc. COVID-19 first cases were observed in Wuhan, China in December 2019, which seems to be the origin of SARS-CoV-2 virus. As of April 2020, there are more than 3.04 million confirmed cases, with 211 thousand deaths globally. Vaccines and commercial detection kits are mostly in the developmental stages to combat this viral infection, and currently, chloroquine and hydroxychloroquine drugs are being used for treatment, but there is no approved drug or vaccine triggering the immune response in the body against SARS-CoV-2 in the market.

In the present study, an integrated bioinformatics approach was used to identify expressed T- and B-cell epitopes from RNA-seq data of SARS-CoV-2 infection in normal human bronchial epithelial (NHBE) and human adeno carcinomic alveolar basal epithelial (A549). To the best of our knowledge, no previous study has been reported a list of expressed T-and B-cell epitopes for multi-epitope based vaccine development. The specific objectives of this research study were: (1) SARS-CoV-2 transcriptome construction and annotation to explore expressed region of SARS-CoV-2 genome, (2) identification of potential T-and B-cell epitopes by using SARS-CoV-2 transcriptome, (3) modelling and docking studies to explore structural compatibility of epitopes with MHC complexes and B-cell receptors, and (4) gene expression of MHC class I and II genes by using RNA-seq data.

## 2. Material and Method

### 2.1. SARS-CoV-2 data retrieval, processing and transcriptome assembly

Due to the recent outbreak of SARS-CoV-2, several countries were started to generate molecular resources to understand pandemic caused by SARS-CoV-2. We used publicly available transcriptome data (PRJNA615032) of SARS-CoV-2 infection in A549 and NHBE cell lines [13]. All the available data were download from the sequence read archive of NCBI database and fastq-dump program of SRAtoolkit [14] was used to extract fastq reads. Quality assessment and control of RNA-seq data was performed through the FastQC version 0.11.5 [15], MultiQC version 1.8 [16] and trimmomatic version 0.39 software [17]. All high-quality reads were mapped over SARS-CoV-2 isolate Wuhan-Hu-1(MN908947.3) by using HISAT2 version 2.1.0 on default parameters [18]. Samtools version 1.1.0 [19] and Bedtools version 2.26.0 [20] were used to extract all the mapped read from each sample and extracted reads were used to construct *de novo* assemblies by using the Trinity assembler version 2.5.1 [21]. TransDecoder program [22] was used to generate protein sequence from assembled transcriptome. Kallisto, a pseudo aligner for bulk RNA-seq data alignment, was used for expression quantification [23].

### 2.2. T-Cell epitope prediction from SARS-CoV-2 transcriptome sequences

T-cell epitopes are short peptide fragments of infectious agents such as viruses and bacteria, which has potential to induce specific immune responses and can be used as a key molecular resource for epitope-based vaccine design. NetCTL 1.2 program [24] was used to predict cytotoxic T-lymphocyte (CTL) epitopes from protein sequences, translated from assembled SARS-CoV-2 transcriptome. NetCTL 1.2 has 12 super types i.e. A1, A2, A3, A24, A26, B7, B8, B27, B39, B44, B58, and B62, and combined score of proteasomal C terminal cleavage for CD8^+^ T-cell epitopes, MHC class I binding, and TAP transport efficiency was considered for epitope identification at threshold value 1.00. To explore antigenic potential of identified peptides, VaxiJen v2.0 [25] server was used at threshold value 0.4, and IEDB program (http://tools.iedb.org/immunogenicity/) was used to identify immunogenicity score for each identified epitopes. Combination of antigenicity and immunogenicity was used to select highly antigenic and immunogenic peptide for further analysis. To explore MHC genes and alleles, all the identified epitopes were analysed through IEDB program mhci (http://tools.iedb.org/mhci/download/) and mhcii (http://tools.iedb.org/mhcii/download/). Peptide length of predicted epitopes 9.0 and inhibitory concentration (IC_50_) value less than or equal to 200nM were selected as parameters for the identification of MHC class I and II binding genes and alleles. Conservation level of selected epitopes were calculated from IEDB program conservancy (http://tools.iedb.org/conservancy/) by using transcriptome sequences, and publically available SARS-CoV-2 protein sequences. IEDB population coverage tool (http://tools.iedb.org/population/) was used to analyze population coverage through predicted MHC alleles of epitopes [26, 27]. Peptide toxicity prediction was performed through the ToxinPred web server [28]. Cross-reactivity with host antigenic proteins might leads to adverse immune responses. Therefore, selected epitopes were checked for similarities with the human proteome sequences (Homo sapiens: GRCh38) through standalone NCBI BLAST similarity search tool.

### 2.3. B-cell epitopes prediction from SARS-CoV-2 transcriptome sequences

Sequence and structure based approaches were used to identify B-cell epitopes. In the sequence based approach, VaxiJen server was used to identify most antigenic proteins from translated transcriptome, and BepiPred-2.0 program [29] was used to identify B-cell epitopes from the identified antigenic proteins. IEDB conformational B-cell prediction tool ElliPro (http://tools.iedb.org/ellipro/) was used to predict epitopes based on protein structure with the parameters PI (protrusion index) value 0.8 as minimum score, and 7Å as maximum distance [30]. Protein sequences of SARS-CoV-2 transcriptome were showed strong sequence similarity with modelled 3D structure of SARS-CoV-2 genome at Zhang lab. Hence, we downloaded 24 structure of SARS-CoV-2 genome from Zhang lab (https://zhanglab.ccmb.med.umich.edu/) for structure-based epitope prediction. Epitopes identified by both the approaches were evaluated for toxicity, antigenicity, and immunogenicity same as done for T-cell epitopes. Cross-reactivity of selected epitopes were checked with human proteome sequences.

### 2.4. Molecular docking studies

Selected epitopes were used for structural compatibility and interaction analysis with available MHC class I and II genes, and B-cell receptor structures. Protein structure of MHC class genes, and B-cell receptors were retrieved from RCSB Protein Data Bank (PDB) (https://www.rcsb.org/) in PDB format. To identify the interactions between predicted T- and B-cell epitopes and protein receptors, the molecular docking studies were performed using AutoDockTools, AutoDock Vina and CABS-dock server [31–33]. Structural compatibility and interaction prediction between peptide and receptor in real world is even more complex and challenging. Therefore, initial screening of peptides for receptors were performed through Autodock, whereas CABSdocks, considered full flexibility of peptide and small fluctuation in receptor backbone, was used for final binding and compatibility analysis. Protein complex visualization and hydrogen bonds were calculated through UCSF chimera [34] and LIGPLOT software package [35], and also ensured the number of genuine hydrogen bonds through cavity prediction by using D3Packets server [36].

## 3. Results

T- and B-cells are the working horses of the adaptive immune system which are capable to produce immunological protection against specific pathogens. The function of T- and B cells are based on the recognition of antigens through specialized receptors and recognized antigenic regions are known as epitopes. Therefore, identification of epitopes for pathogens is crucial for the understanding of disease etiology, disease diagnostics, and epitope-based vaccine development. In this study, an effort was made to identify expressed T- and B-cell epitopes from RNA-seq data of SARS-CoV-2 infection in human NHBE and A549 cell lines. To achieve this goal, various bioinformatics approaches, tools and software were used as summarized in figure-1.

**Figure-1.**
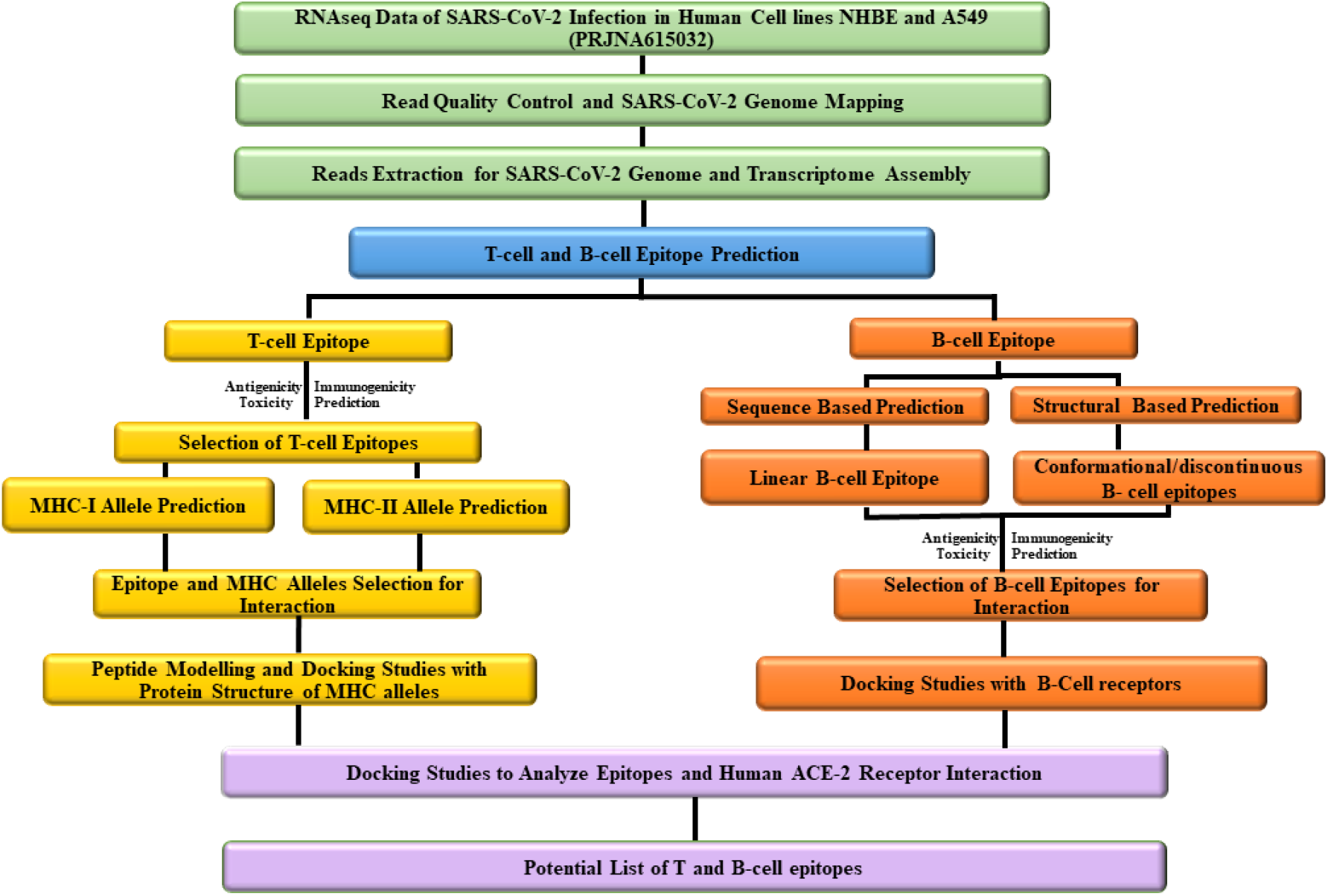
Schematic representation of used approach for transcriptome assembly from RNA-seq data of SARS-CoV-2 infection in human cell lines, epitope identification, and molecular docking studies.

### 3.1 SARS-CoV-2 transcriptome assembly and annotation

*De novo* SARS-CoV-2 transcriptome was constructed by using publicly available transcriptome data from SRA project (PRJNA615032). Transcriptome data was generated to the study of SARS-CoV-2 infection in human cell lines NHBE and A549 [13]. Extracted reads were trimmed by removing adapter and low quality sequences by using trimmomatic-0.39. Reads with a length of less than 20 bps were also removed from dataset [37]. In order to develop SARS-CoV-2 transcriptome assembly, raw reads were aligned to the SARS-CoV-2 genome (Accession number: MN908947.3, Wuhan-Hu-1 isolate) by using the RNA-seq alignment tool HISAT2 on default conditions. After read mapping, Samtools and Bedtools were used to extract mapped RNA reads on SARS-CoV-2 genome. Total, 87,716 reads were extracted from all the samples. Detail description of experiment, sample name, description, and number of mapped reads per sample over SARS-CoV-2 genome were given in supplementary material file1 (Table – S1). All the extracted RNA-seq reads were used to construct *de novo* SARS-CoV-2 transcriptome through Trinity software. In total, 54,814 bases were assembled into 27 transcripts with median contig length 650 bps, N50 value of 10,677 bps and approximate average transcript length of 2030 bps. The generated transcriptome assembly was clustered at 90 % sequence identity through CD-HIT software that produced 27 non-redundant transcripts, the same number of non-redundant transcripts showed the sequence variability among assembled transcripts. Non-redundant transcripts were translated into 44 protein sequences through TransDecoder program. Protein sequences were annotated against Uniport databases by using BLAST similarity search at evalue threshold 1e-10 (Table – 1).

**Table-1.**
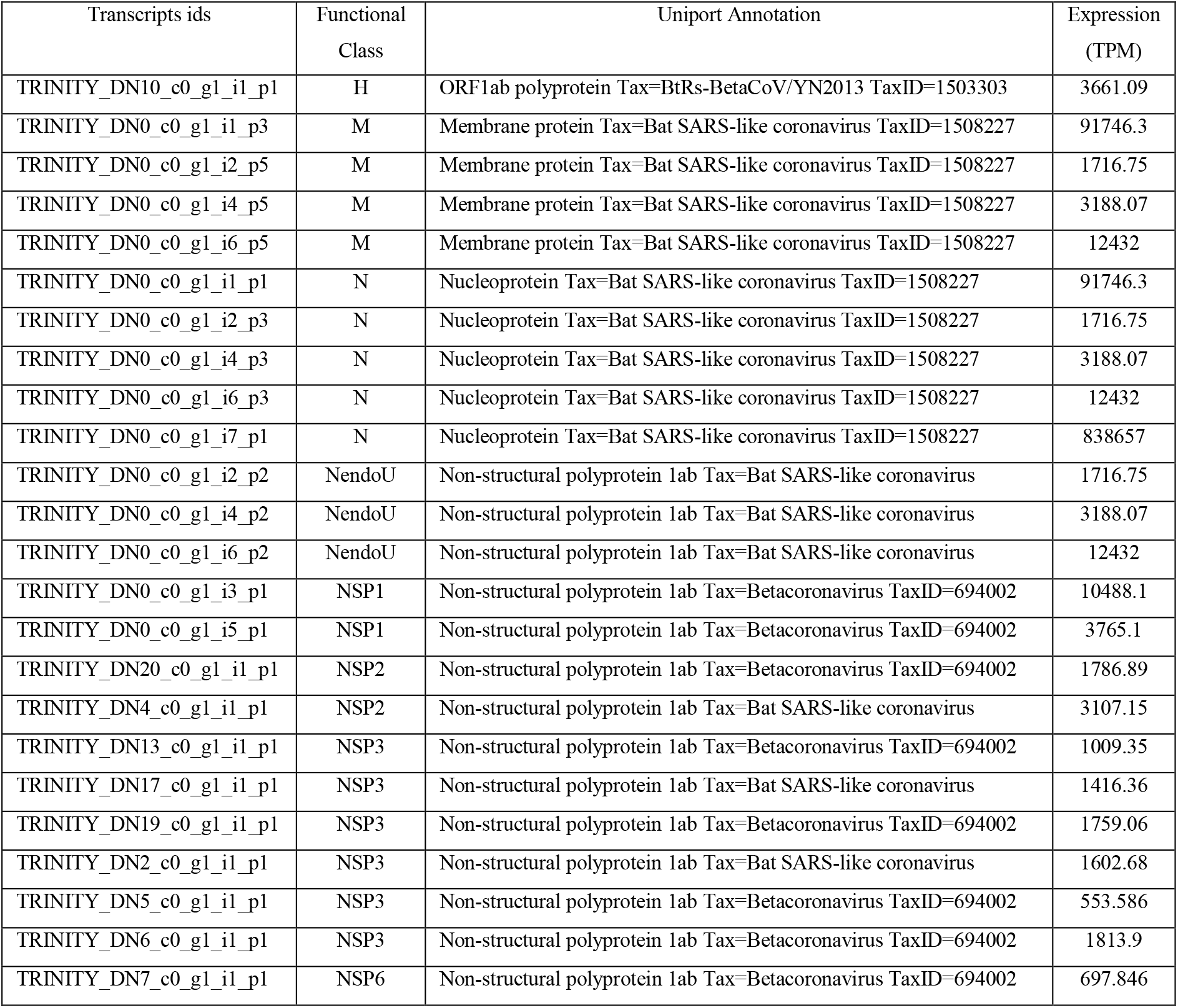
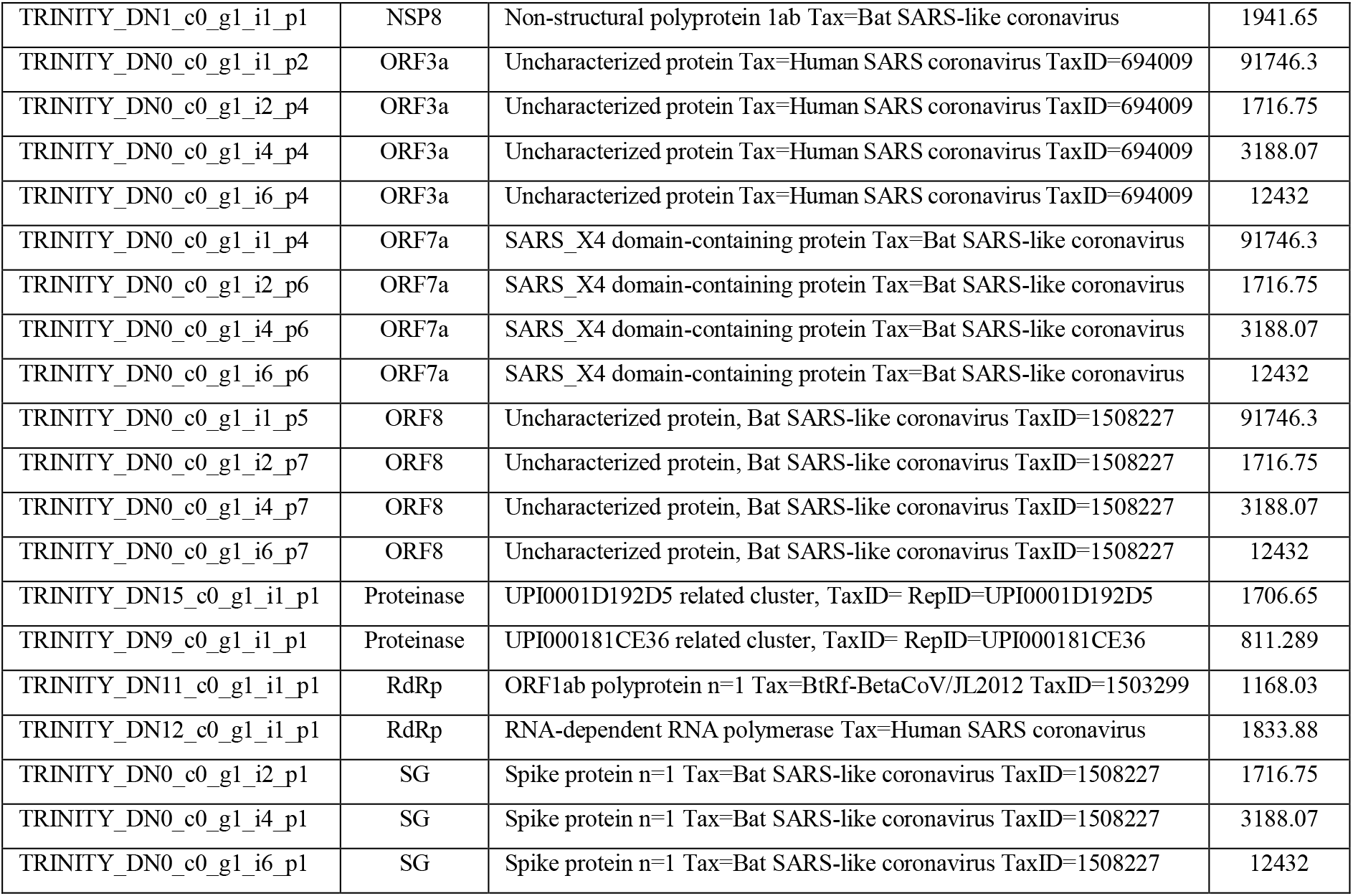
SARS-CoV-2 transcriptome annotation along with expression values. H: Helicase, M: Membrane protein, N: Nucleoprotein, NendoU: Uridylate-specific endoribonuclease, NSP: Non-structural protein, ORF: open reading frame, SG: Surface glycoprotein, RdRp: RNA-dependent RNA polymerase, TPM: Transcript per million.

| Transcripts ids | Functional Class | Uniport Annotation | Expression (TPM) |
| --- | --- | --- | --- |
| TRINITY_DN10_c0_g1_i1_p1 | H | ORF1ab polyprotein Tax=BtRs-BetaCoV/YN2013 TaxID=1503303 | 3661.09 |
| TRINITY_DN0_c0_g1_i1_p3 | M | Membrane protein Tax=Bat SARS-like coronavirus TaxID=1508227 | 91746.3 |
| TRINITY_DN0_c0_g1_i2_p5 | M | Membrane protein Tax=Bat SARS-like coronavirus TaxID=1508227 | 1716.75 |
| TRINITY_DN0_c0_g1_i4_p5 | M | Membrane protein Tax=Bat SARS-like coronavirus TaxID=1508227 | 3188.07 |
| TRINITY_DN0_c0_g1_i6_p5 | M | Membrane protein Tax=Bat SARS-like coronavirus TaxID=1508227 | 12432 |
| TRINITY_DN0_c0_g1_i1_p1 | N | Nucleoprotein Tax=Bat SARS-like coronavirus TaxID=1508227 | 91746.3 |
| TRINITY_DN0_c0_g1_i2_p3 | N | Nucleoprotein Tax=Bat SARS-like coronavirus TaxID=1508227 | 1716.75 |
| TRINITY_DN0_c0_g1_i4_p3 | N | Nucleoprotein Tax=Bat SARS-like coronavirus TaxID=1508227 | 3188.07 |
| TRINITY_DN0_c0_g1_i6_p3 | N | Nucleoprotein Tax=Bat SARS-like coronavirus TaxID=1508227 | 12432 |
| TRINITY_DN0_c0_g1_i7_p1 | N | Nucleoprotein Tax=Bat SARS-like coronavirus TaxID=1508227 | 838657 |
| TRINITY_DN0_c0_g1_i2_p2 | NendoU | Non-structural polyprotein 1ab Tax=Bat SARS-like coronavirus | 1716.75 |
| TRINITY_DN0_c0_g1_i4_p2 | NendoU | Non-structural polyprotein 1ab Tax=Bat SARS-like coronavirus | 3188.07 |
| TRINITY_DN0_c0_g1_i6_p2 | NendoU | Non-structural polyprotein 1ab Tax=Bat SARS-like coronavirus | 12432 |
| TRINITY_DN0_c0_g1_i3_p1 | NSP1 | Non-structural polyprotein 1ab Tax=Betacoronavirus TaxID=694002 | 10488.1 |
| TRINITY_DN0_c0_g1_i5_p1 | NSP1 | Non-structural polyprotein 1ab Tax=Betacoronavirus TaxID=694002 | 3765.1 |
| TRINITY_DN20_c0_g1_i1_p1 | NSP2 | Non-structural polyprotein 1ab Tax=Betacoronavirus TaxID=694002 | 1786.89 |
| TRINITY_DN4_c0_g1_i1_p1 | NSP2 | Non-structural polyprotein 1ab Tax=Bat SARS-like coronavirus | 3107.15 |
| TRINITY_DN13_c0_g1_i1_p1 | NSP3 | Non-structural polyprotein 1ab Tax=Betacoronavirus TaxID=694002 | 1009.35 |
| TRINITY_DN17_c0_g1_i1_p1 | NSP3 | Non-structural polyprotein 1ab Tax=Bat SARS-like coronavirus | 1416.36 |
| TRINITY_DN19_c0_g1_i1_p1 | NSP3 | Non-structural polyprotein 1ab Tax=Betacoronavirus TaxID=694002 | 1759.06 |
| TRINITY_DN2_c0_g1_i1_p1 | NSP3 | Non-structural polyprotein 1ab Tax=Bat SARS-like coronavirus | 1602.68 |
| TRINITY_DN5_c0_g1_i1_p1 | NSP3 | Non-structural polyprotein 1ab Tax=Betacoronavirus TaxID=694002 | 553.586 |
| TRINITY_DN6_c0_g1_i1_p1 | NSP3 | Non-structural polyprotein 1ab Tax=Betacoronavirus TaxID=694002 | 1813.9 |
| TRINITY_DN7_c0_g1_i1_p1 | NSP6 | Non-structural polyprotein 1ab Tax=Betacoronavirus TaxID=694002 | 697.846 |
| TRINITY_DN1_c0_g1_i1_p1 | NSP8 | Non-structural polyprotein 1ab Tax=Bat SARS-like coronavirus | 1941.65 |
| TRINITY_DN0_c0_g1_i1_p2 | ORF3a | Uncharacterized protein Tax=Human SARS coronavirus TaxID=694009 | 91746.3 |
| TRINITY_DN0_c0_g1_i2_p4 | ORF3a | Uncharacterized protein Tax=Human SARS coronavirus TaxID=694009 | 1716.75 |
| TRINITY_DN0_c0_g1_i4_p4 | ORF3a | Uncharacterized protein Tax=Human SARS coronavirus TaxID=694009 | 3188.07 |
| TRINITY_DN0_c0_g1_i6_p4 | ORF3a | Uncharacterized protein Tax=Human SARS coronavirus TaxID=694009 | 12432 |
| TRINITY_DN0_c0_g1_i1_p4 | ORF7a | SARS_X4 domain-containing protein Tax=Bat SARS-like coronavirus | 91746.3 |
| TRINITY_DN0_c0_g1_i2_p6 | ORF7a | SARS_X4 domain-containing protein Tax=Bat SARS-like coronavirus | 1716.75 |
| TRINITY_DN0_c0_g1_i4_p6 | ORF7a | SARS_X4 domain-containing protein Tax=Bat SARS-like coronavirus | 3188.07 |
| TRINITY_DN0_c0_g1_i6_p6 | ORF7a | SARS_X4 domain-containing protein Tax=Bat SARS-like coronavirus | 12432 |
| TRINITY_DN0_c0_g1_i1_p5 | ORF8 | Uncharacterized protein, Bat SARS-like coronavirus TaxID=1508227 | 91746.3 |
| TRINITY_DN0_c0_g1_i2_p7 | ORF8 | Uncharacterized protein, Bat SARS-like coronavirus TaxID=1508227 | 1716.75 |
| TRINITY_DN0_c0_g1_i4_p7 | ORF8 | Uncharacterized protein, Bat SARS-like coronavirus TaxID=1508227 | 3188.07 |
| TRINITY_DN0_c0_g1_i6_p7 | ORF8 | Uncharacterized protein, Bat SARS-like coronavirus TaxID=1508227 | 12432 |
| TRINITY_DN15_c0_g1_i1_p1 | Proteinase | UPI0001D192D5 related cluster, TaxID= RepID=UPI0001D192D5 | 1706.65 |
| TRINITY_DN9_c0_g1_i1_p1 | Proteinase | UPI000181CE36 related cluster, TaxID= RepID=UPI000181CE36 | 811.289 |
| TRINITY_DN11_c0_g1_i1_p1 | RdRp | ORF1ab polyprotein n=1 Tax=BtRf-BetaCoV/JL2012 TaxID=1503299 | 1168.03 |
| TRINITY_DN12_c0_g1_i1_p1 | RdRp | RNA-dependent RNA polymerase Tax=Human SARS coronavirus | 1833.88 |
| TRINITY_DN0_c0_g1_i2_p1 | SG | Spike protein n=1 Tax=Bat SARS-like coronavirus TaxID=1508227 | 1716.75 |
| TRINITY_DN0_c0_g1_i4_p1 | SG | Spike protein n=1 Tax=Bat SARS-like coronavirus TaxID=1508227 | 3188.07 |
| TRINITY_DN0_c0_g1_i6_p1 | SG | Spike protein n=1 Tax=Bat SARS-like coronavirus TaxID=1508227 | 12432 |

### 3.2. T-cell epitopes identification of SARS-CoV-2 transcriptome

T-cell epitopes are presented by MHC class I and II that are recognized by two distinct subsets of T-cells, CD8^+^ and CD4^+^ T-cells, respectively. NetCTL 1.2 program was used for the prediction T-cell epitopes, and 1144 epitopes were selected at combined prediction threshold value 1.0 for 12 super type categories i.e. A1 (330), A24(314), A26(242), A2(247), A3(328), B27(175), B39(263), B44(133), B58(284), B62(473), B7(157), and B8(193). The predicted T-cell epitopes were further evaluated for antigenicity by VaxiJen server and immunogenicity by IEDB prediction tools, and 598 and 625 epitopes were shown antigenicity and immunogenicity potential respectively. Finally, 598 antigenicity and immunogenicity T-cell epitopes were selected for further study. To determine epitopes potential to elicit effective immune response, 598 selected epitopes were used to explore interacting MHC alleles through IEDB program. Peptide length nine and the IC_50_ value = < 200nM were selected as parameters for the identification of MHC class I binding gene and alleles. MHC class I alleles such as HLA-A, HLA-B, and HLA-C were recognized with parameter human as MHC source species and IEBD recommended method to predict a distinct set of MHC class I alleles for all selected 598 epitopes. In MHC-I allele analysis, HLA-A type alleles were found as more frequently occurring alleles (HLA-A*01:01, HLA-A*02:01, HLA-A*02:03, HLA-A*02:06, HLA-A*03:01, HLA-A*11:01, HLA-A*23:01, HLA-A*24:02, HLA-A*26:01, HLA-A*30:01, HLA-A*30:02, HLA-A*31:01, HLA-A*32:01, HLA-A*33:01, HLA-A*68:01, HLA-A*68:02) than HLA-B type alleles (HLA-B*07:02, HLA-B*08:01, HLA-B*15:01, HLA-B*35:01, HLA-B*40:01, HLA-B*44:02, HLA-B*44:03, HLA-B*53:01, HLA-B*57:01). In our analysis, HLA-C class genes were not found for any epitopes. ToxinPred was used to explore toxicity of predicted epitopes. Finally, 40 CD8^+^ T-cell epitopes (Table-2) were selected as high immunogenic, antigenic, non-toxic epitopes with good binding affinity to MHC class I alleles. Further, these epitopes were explored for SARS-CoV-2 proteome for functional characterization.

**Table – 2.**
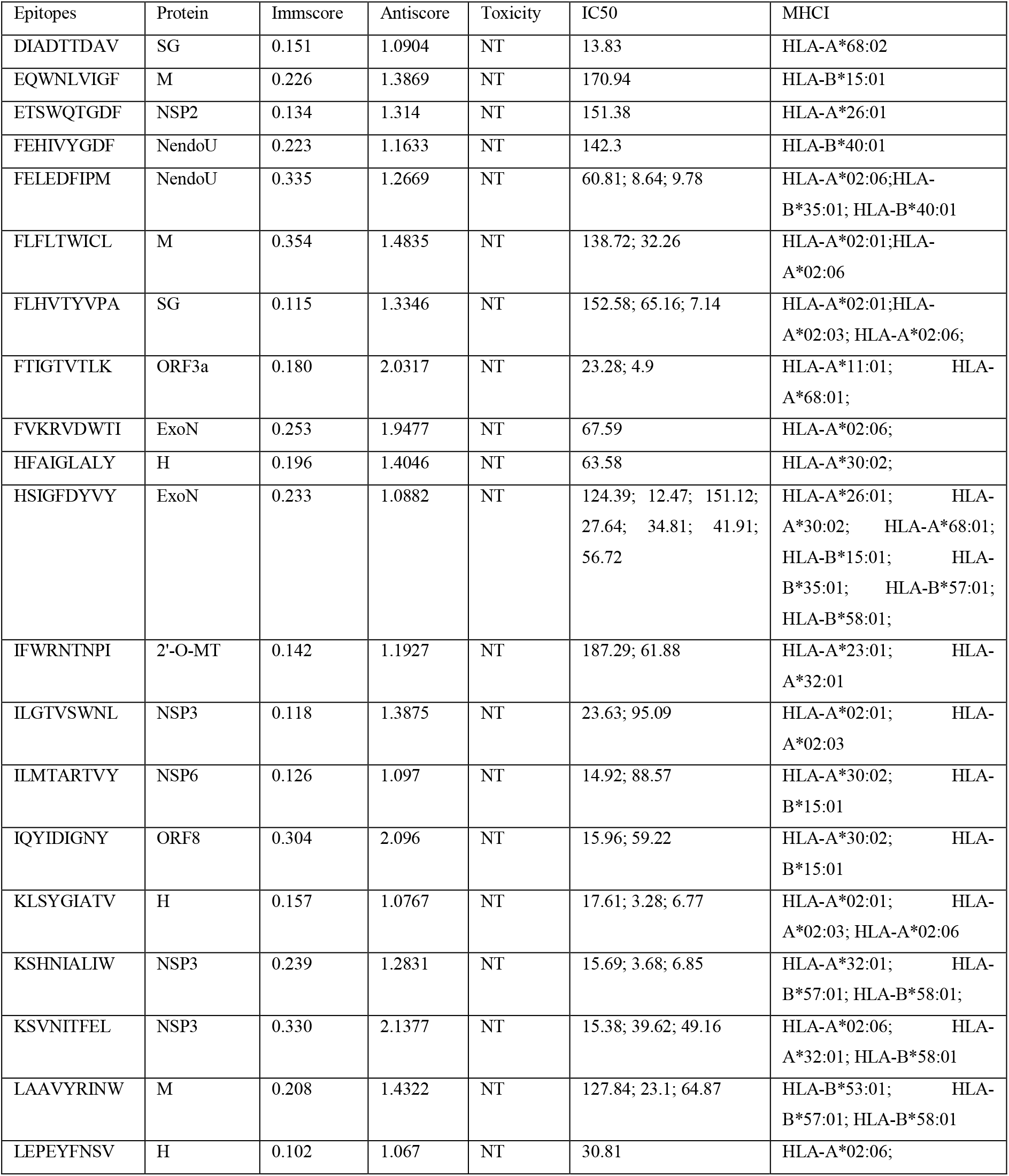
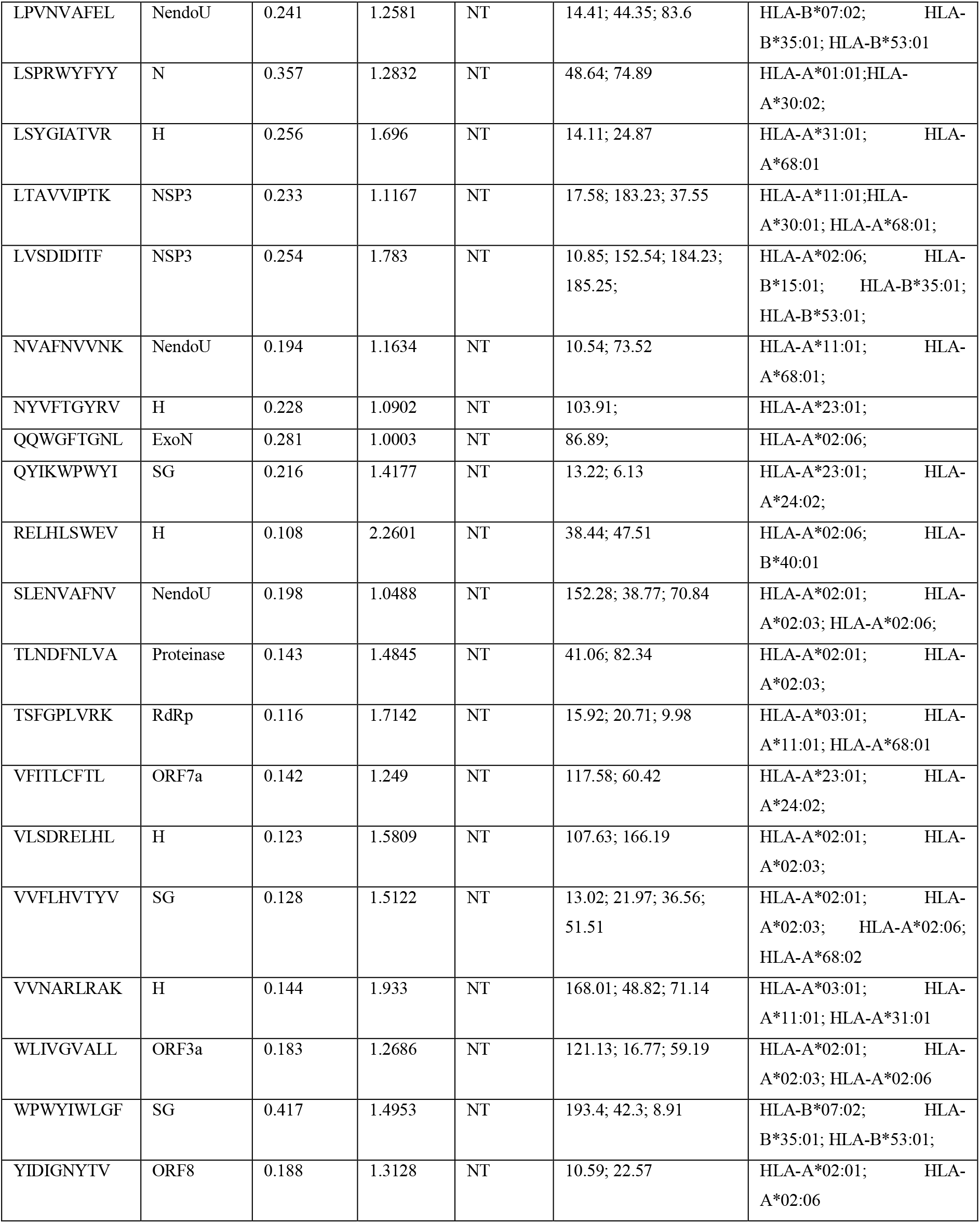
List of potential T-cell epitopes for MHC class I. Immscore: Immunogencity score, Antiscore: Antigencity score, IC: inhibitory constant, NT: Non-toxic, H: Helicase, M: Membrane protein, N: Nucleoprotein, NendoU: Uridylate-specific endoribonuclease, NSP: Non-structural protein, ORF: open reading frame, SG: Surface glycoprotein, RdRp: RNA-dependent RNA polymerase, 2’-O-MT: 2’-O-methyltransferase, ExoN: Guanine-N7 methyltransferase

Similarly, MHC class II gene and allele’s prediction was performed through IEDB analysis resources by using the same parameters as for MHC class I except the selection of SMM method for the prediction of a distinct set of MHC class II alleles. Total, 4072 epitopes were identified from protein sequences with good binding affinity to MHC class II alleles. To select MHC class II alleles and epitopes, we decided to take those MHC class II alleles and epitopes which have MHC class I epitopes as a core sequence. Total, 34 MHC class II epitope sequences (15-mer) were selected by using previously selected 40 (9-mer) antigenic and immunogenic epitope sequences as core sequences (Supplementary file1: Table-S2). Among all MHC-II alleles, HLA-DPA1*01:03/DPB1*04:01, HLA-DPA1*02:01/DPB1*14:01, HLA-DRB3*02:02, HLA-DRB1*01:01, HLA-DRB1*07:01, HLA-DPA1*01:03/DPB1*02:01, and HLA-DRB5*01:01were the most abundant alleles. Among all, various 9-mer epitopes such as LSPRWYFYY, KSVNITFEL, IQYIDIGNY, EQWNLVIGF, DIADTTDAV, TSFGPLVRK and RELHLSWEV were also have core sequence among 15-mer MHC class II alleles epitopes. But IC_50_ value of these epitopes were more than 200nM. After conservation analysis, twelve most antigenic and immunogenic MHC class II epitopes (APHGVVFLHVTYVPA, FLHVTYVPAQEKNFT, GVVFLHVTYVPAQEK, HGVVFLHVTYVPAQE, PHGVVFLHVTYVPAQ, QSAPHGVVFLHVTYV, QYIKWPWYIWLGFIA, SAPHGVVFLHVTYVP, VFLHVTYVPAQEKNF, VVFLHVTYVPAQEKN, and YIKWPWYIWLGFIAG) were selected which contains 9-mer core sequences of four epitopes from previously selected 40 epitopes. Peptide sequences of 9-mer epitopes, CD4^+^ T-cell epitopes sequence and MHC class II alleles were given in Table-3.

**Table – 3:**
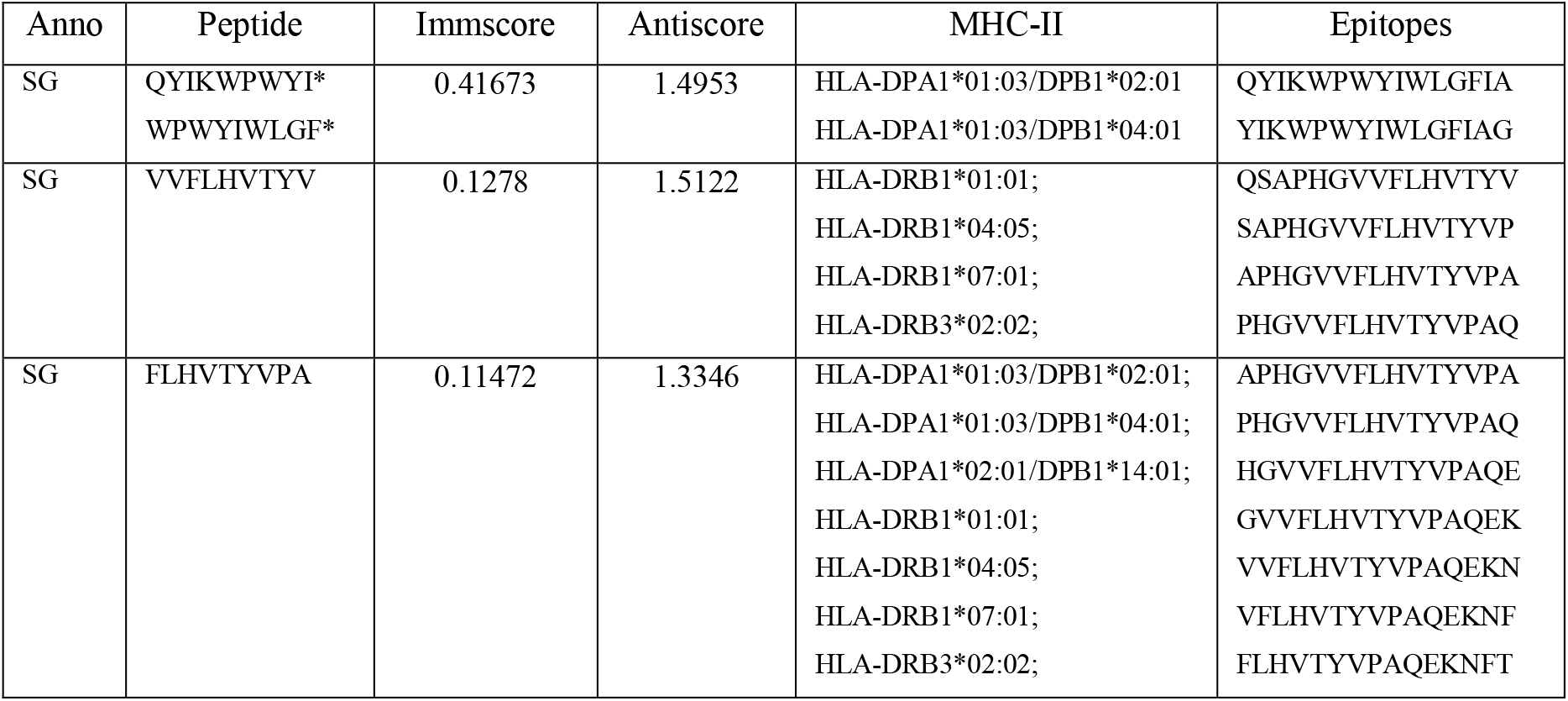
List of potential non-toxic, conserved epitopes for MHC class II along with their core 9-mer epitopes. Anno: Annotation, Immscore: Immunogencity score, Antiscore: Antigencity score, SG: Surface glycoprotein, *: overlapping epitopes

### 3.3. Population coverage analysis

MHC molecules can form complexes with millions of epitopes which are reflecting the polymorphic nature of MHC genes. If MHC polymorphism occurs in peptide-binding region, binding specificity of MHC molecules will be changed. MHC variability has evolutionary advantage to identify variety of pathogens. But genetic variability among MHC alleles are also a major obstacles in the development of peptide-based vaccines. Therefore, population coverage is an important criterion to design a generalized an effective vaccine[38].

In population coverage analysis, MHC class I allele’s of 40 epitopes and MHC class II allele’s of 33 epitopes were used, and a significant population coverage was found for different geographic regions around the world (Figure-2). MHC class alleles of selected epitopes were covered approximately 90% of the world population. Highest population coverage was found for Sweden (100%) which was closely followed by England, Germany, France, Belgium, United States, Russia, Italy, South Korea, Japan, Mexico, Iran, Chile, Brazil, China, Singapore, Pakistan, India, Spain, Thailand, Israel, Philippines, Australia, and Vietnam with a population coverage of 99.99%, 99.99%, 99.97%, 99.87%, 99.87%, 99.81%, 99.78%, 99.66%, 99.55%, 99.02%, 98.84%, 98.55%, 98.43%, 97.6%, 97.24%, 97.13%, 97.05%, 96.9%, 96.48%, 96.47%, 96.41%, 96.17%, and 95.91% respectively. The lowest population coverage were found for Canada (38.31%), Srilanka (42.04) and Ukrain (46.48). United States and Europe has highest number of COVID-19 cases [39]. Hence, the population coverage prediction is essential for vaccine design. Population of ethnic groups were also significantly covered (Supplementary file1: Table – S3), and average coverage for ethnic group across the world is around 93%.

**Figure – 2.**
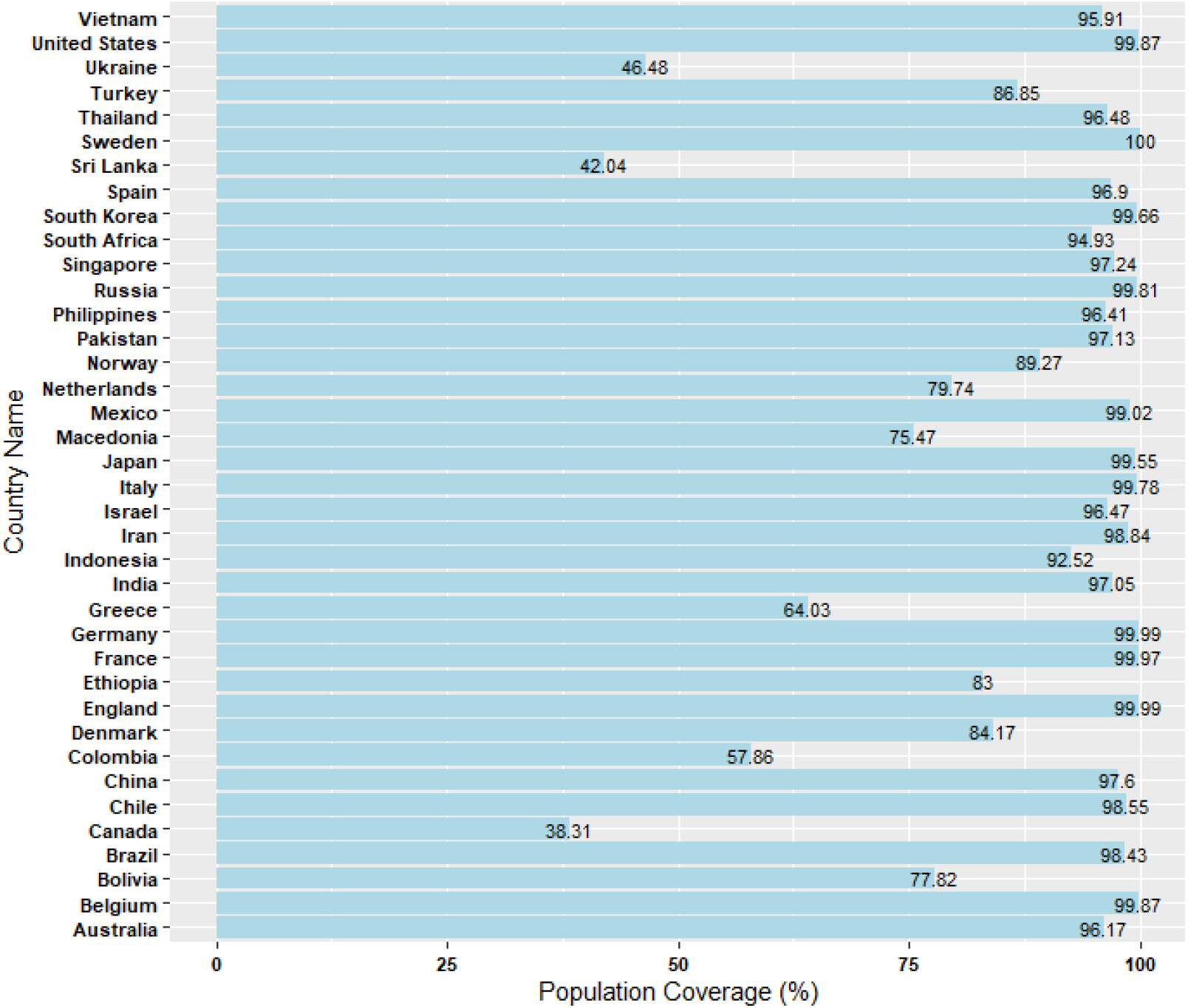
Global population coverage through identified epitopes.

### 3.4. B-cell epitope identification of SARS-CoV-2 transcriptome

B-cell epitope is a precise region of the antigenic protein that is detected by B-cell receptors (BCR) through membrane-bound immunoglobulins. Once B-cell activated, it secretes soluble forms of the immunoglobulins to neutralize antigenic proteins. Thus, B-cell epitope and B-cell receptor information is essential for epitope-based vaccine design. Prediction of B-cell epitopes was performed by using protein sequences of the assembled transcriptome and SARS-CoV-2 protein structures. Total, 330 B-cells epitopes were predicted through BepiPred-2.0 program by using protein sequences, whereas 77 B-cell epitopes were predicted through the IEDB conformational tool ElliPro by using 24 SARS-CoV-2 protein structure, download from Zhang lab. B-cell epitopes prediction from the protein structure is highly useful information for epitope-based vaccine design and development [40, 41]. Therefore, a separate analysis was performed for B-cell epitopes identified from protein structures, and 19 epitopes were identified after toxicity (non-toxic), immunogenicity and antigenicity analysis (Supplementary file1: Table-S4). In order to explore most suitable B-cell epitopes, epitopes generated by both the approaches were combined, and total 130 unique B-cell epitopes were identified from length range 9 to 15. 73 B-cell epitopes were predicted as antigenic epitopes through VaxiJen v2.0 webserver and 53 B-cell epitopes were identified as immunogenic. Total, 16 non-toxic B-cell epitopes were identified with immunogenicity score and antigenicity score more than 0.1 and 0.4 respectively.

Cross-reactivity analysis of epitopes were performed against human proteome sequences through BLAST similarity search, and found that two B-cell epitopes (DNNFCGPDGYPLE,NQDLNGNWYD) showed significant similarity with human proteins ENSP00000390696.1 and ENSP00000263390.3 respectively (Supplementary file1: Table-S5). Finally, 14 B-cell epitope were found for further analysis (Table – 4). On the basis of immunological parameters, the five B-cell epitopes, NSP2 (SEQLDFIDTKRGV, HCGETSWQTGDFV), Helicase (KGTLEPEYF), NSP3 Papain-like (KTVGELGDVRE), and Surface glycoprotein (LTGTGVLTESNK) were selected from Table – 4 for molecular docking studies of B-cell receptors.

**Table – 4:**
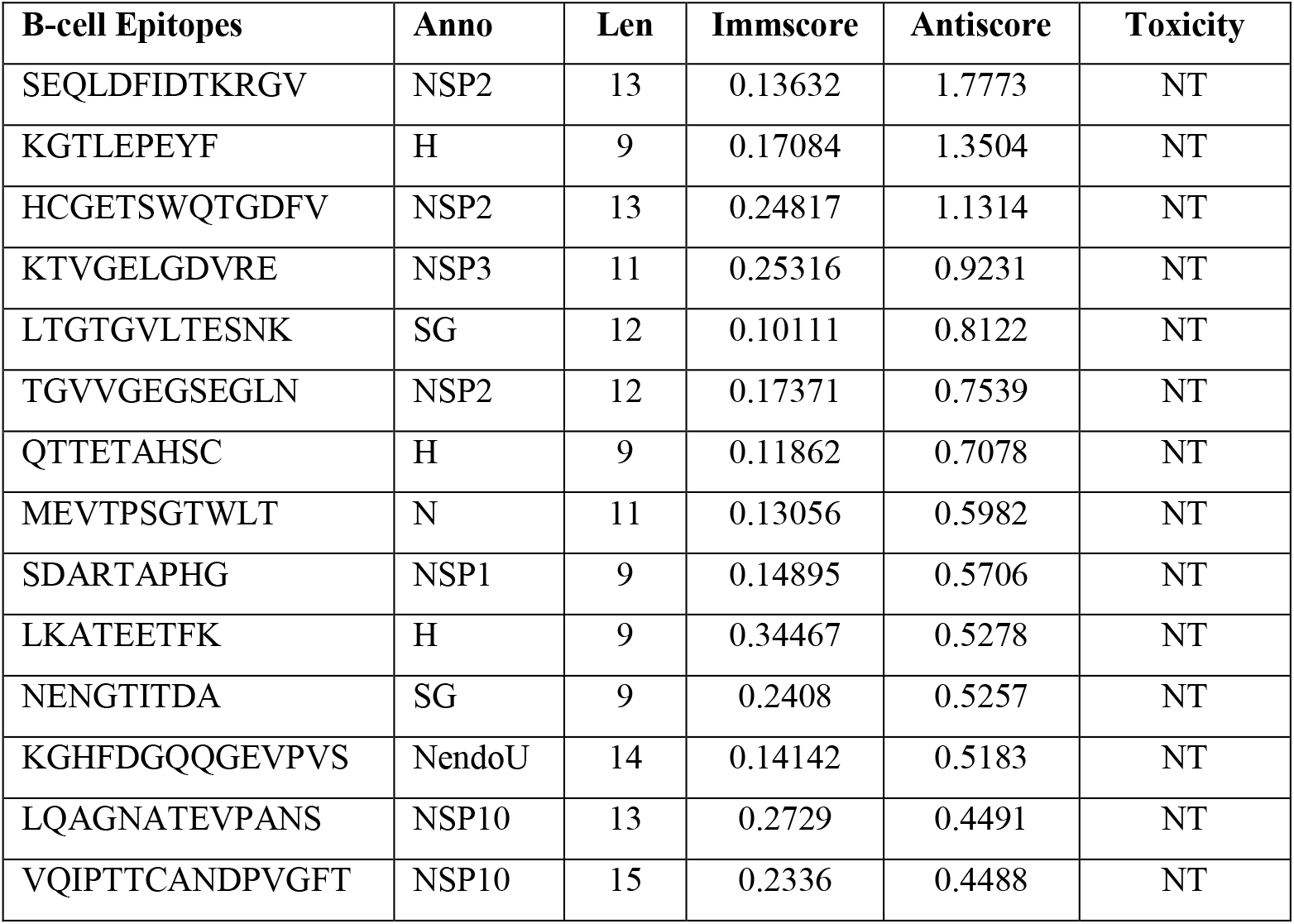
List of B-cell epitopes identified form protein sequences of assembled transcriptome and modelled protein structure of various coding protein of SAR-CoV-2 genome. Immscore: Immunogencity score, Antiscore: Antigencity score, NT: Non-toxic, H: Helicase, M: Membrane protein, N: Nucleoprotein, NendoU: Uridylate-specific endoribonuclease, NSP: Non-structural protein, ORF: open reading frame, SG: Surface glycoprotein, RdRp: RNA-dependent RNA polymerase, 2’-O- MT: 2’-O-methyltransferase, ExoN: Guanine-N7 methyltransferase

### 3.5. Molecular docking analysis

Cellular immunity gets activated when MHC molecules binds to intracellular and extracellular proteins displayed on the cell surface. Structural analysis of epitopes and MHC molecules can improve our knowledge about T-cell based mechanism to reduce disease burden. To explore structural compatibility between T-cell epitopes and MHC complexes, molecular docking studies were performed to analyze binding affinities between MHC complexes and T-cell epitopes. Twenty-two MHC proteins were explored with selected three T-cell epitopes, and the best interaction was identified with the highest binding affinity. The compatible structural model of epitopes (WPWYIWLGF, VVFLHVTYV, and FLHVTYVPA) and the MHC molecules were retaining a binding affinity range from of −136.54 to – 7.12 kcal/mol. Detail description of molecular interaction analysis of peptide VVFLHVTYV and identified protein structure of MHC genes were given in Table – 5. Detail docking descriptions of other two epitopes with MHC molecule were given in supplementary material file1 (Table – S6).

**Table – 5.**
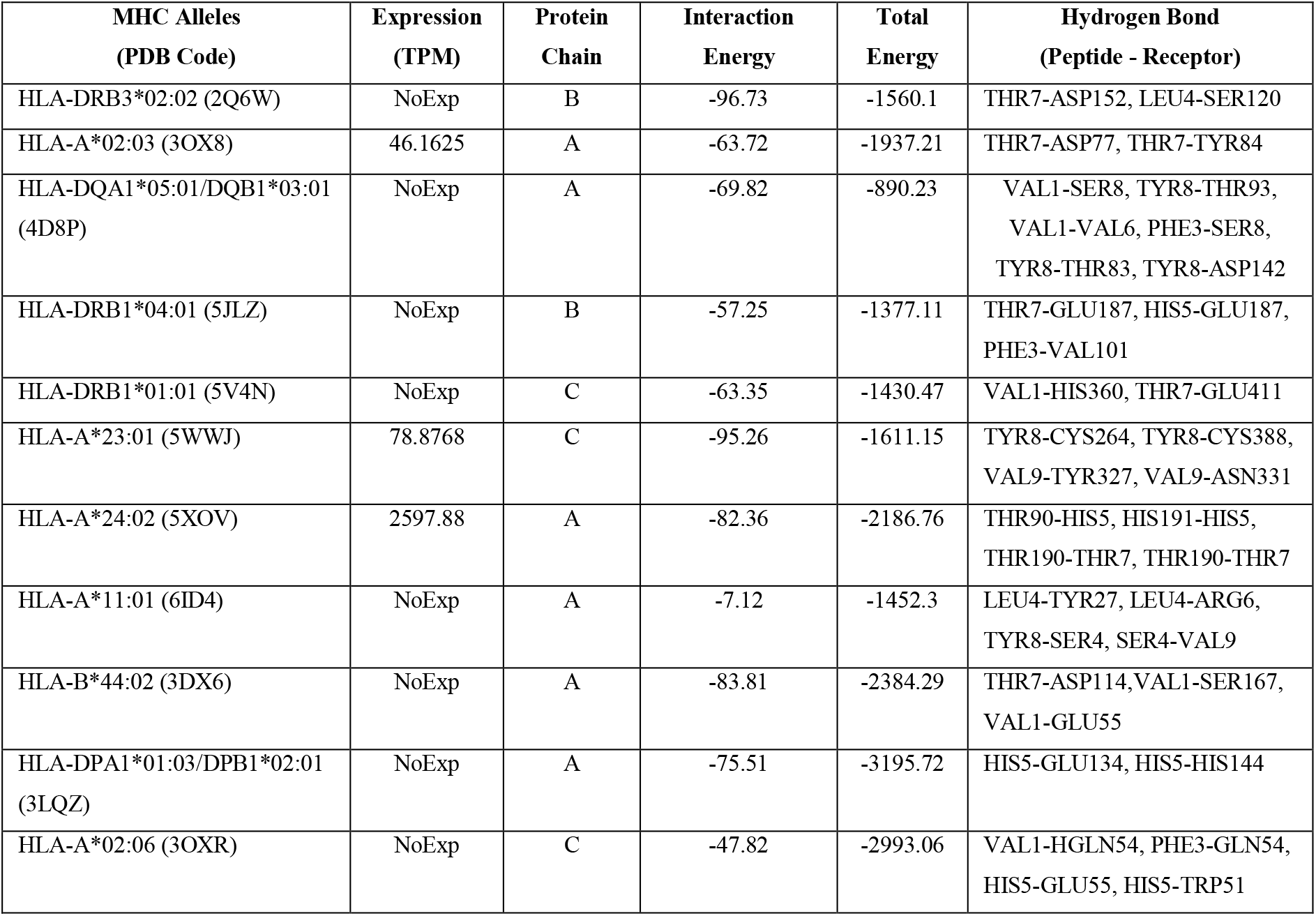
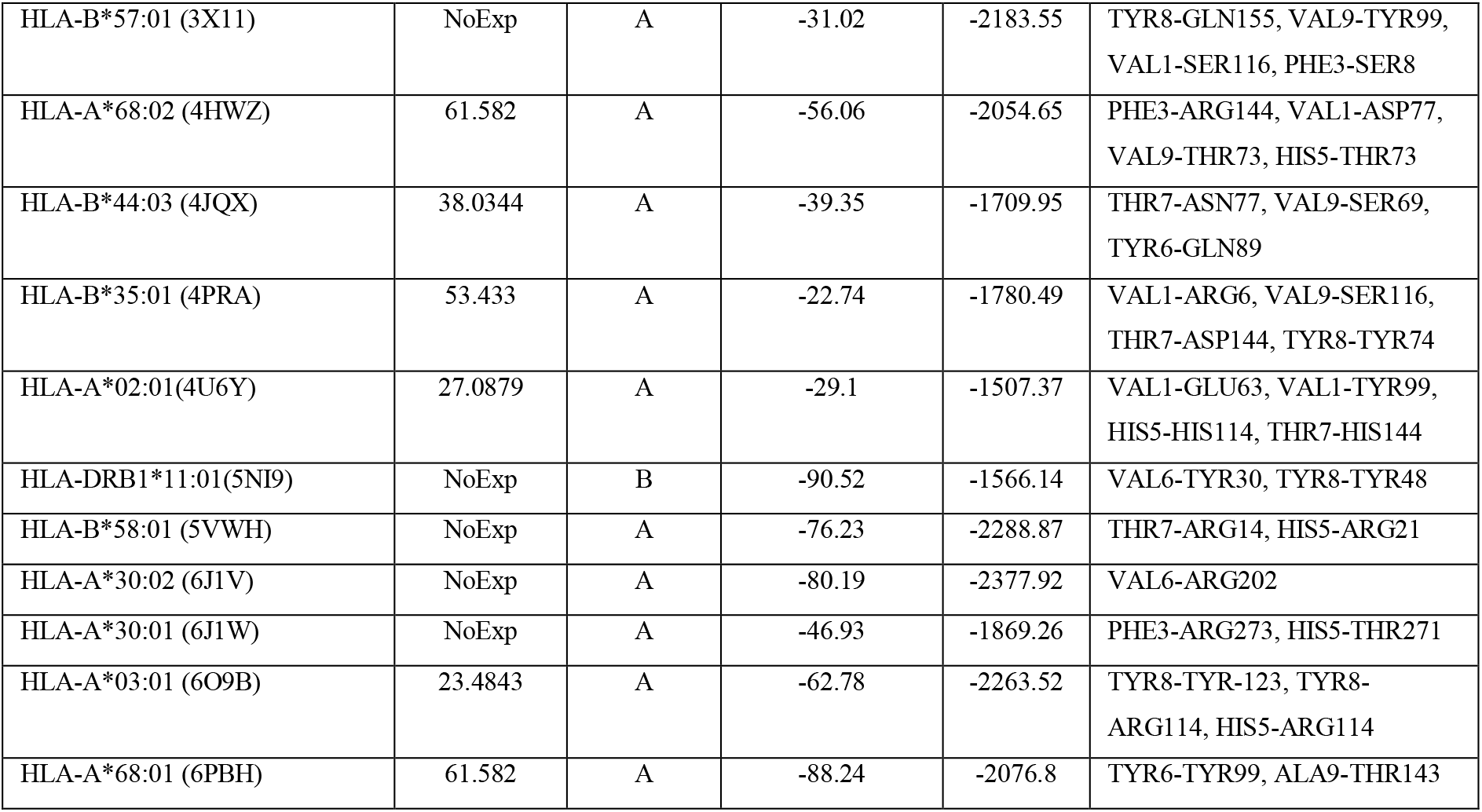
Molecular interaction and docking analysis of identified antigenic and immunogenic T-cell epitope (VVFLHVTYV) and selected protein structures of MHC alleles. TPM (Transcript per million)

MHC class I and II gene expression analysis was performed by using RNA-seq data of cell lines through pseudo aligner Kallisto[23]. Expression values of expressed MHC class alleles were given in supplementary material file1 (Table – S7). HLA-A*24:02 allele was highly expressed among most frequent occurring MHC alleles, but highest interaction of peptide (VVFLHVTYV) was shown with HLA-DRB3*02:02 allele with two hydrogen bonds (THR7-ASP152, LEU4-SER120). The epitope position in protein structure and the binding interactions had shown in Figure-3. We were also ensured genuine hydrogen bonds interaction with receptors through cavity prediction (Supplementary material file2)

**Figure – 3.**
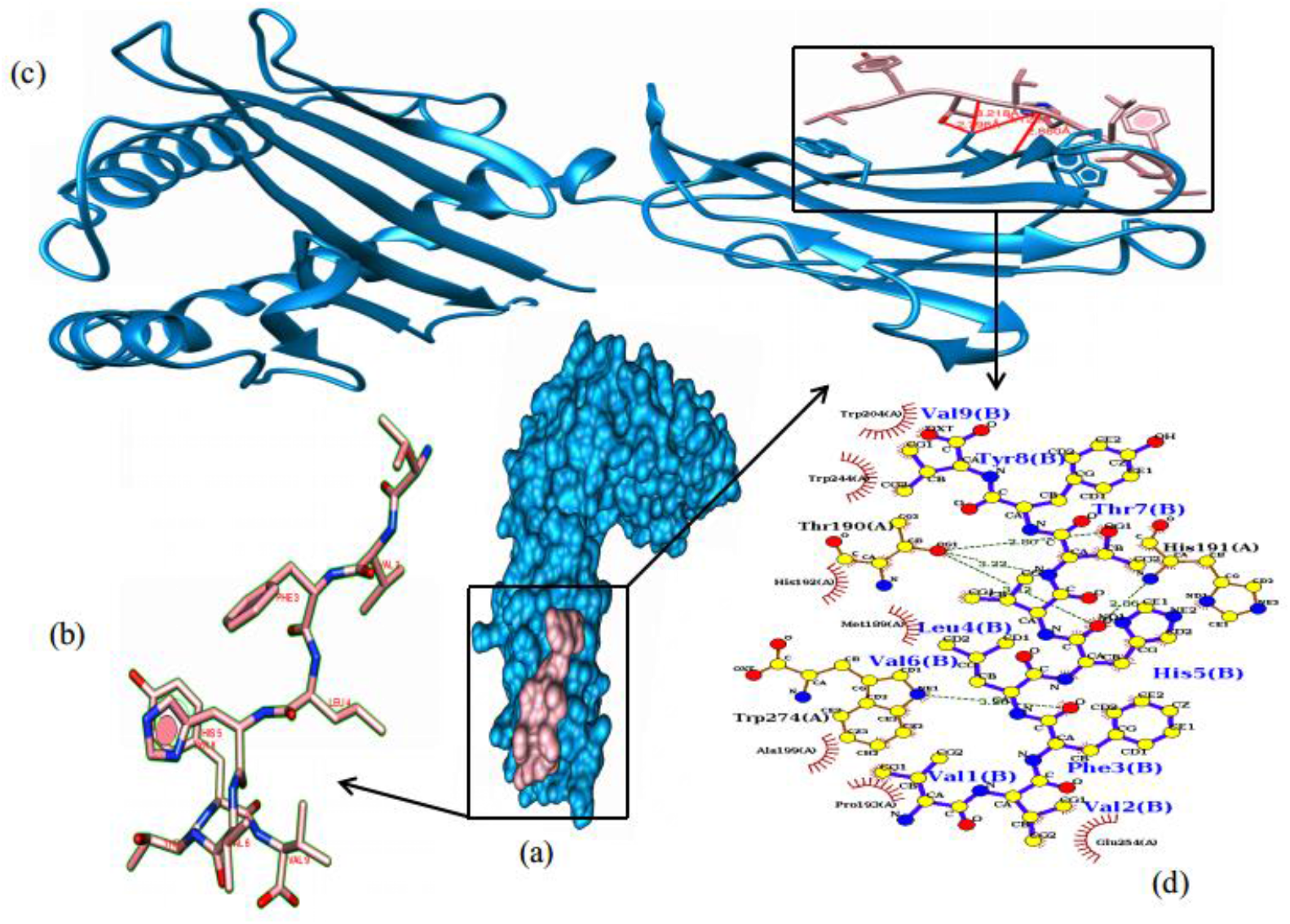
Molecular interaction of peptide (VVFLHVTYV) and protein structure (5XOV) of HLA allele (HLA-A*24:02). a and c) cartoon structure of peptide and class-I HLA allele. b) interaction of peptide and MHC-II alleles (Hydrogen bonds in red colour). d) molecular level description of interaction through LIGPlot software (Hydrogen bonds in green).

Similarly, five most antigenic and immunogenic B-cell epitopes were selected for molecular docking studies with two B-cell receptors (5DRW and 1K1F) through CABSDocks server. 5DRW protein structure is a crystal structure of BCR Fab fragment from subset of chronic lymphocytic leukaemia whereas 1K1F is dimer structure of Bcr-Abl oncoprotein. 1K1F protein structure provided a base to design an inhibitor to disrupt Bcr-Abl oligomerization. Moreover, 1K1F was without Fab region unlike to 5DRW [42]. In our analysis, most of the peptides were shown higher binding affinities with 5DRW than 1K1F. Detail description of docking studies of B-cell receptor and peptides were given in Table – 6.

**Table – 6:** Molecular docking and interaction analysis of B-cell epitope and B-cell protein receptors.

| Peptide | PDB Code | Protein Chain | Interaction Energy | Total Energy | Hydrogen Bonds (Peptide - Receptor) |
| --- | --- | --- | --- | --- | --- |
| SEQLDFIDTKRGV | 5DRW | B | -94.17 | -1656.88 | THR9-PRO49, THR9-SER48, THR9-ARG51, GLU7-LYS55 |
|  | 1K1F | A | -25.53 | -362.98 | THR9-ARG50, TRP7-ARG51, GLU4-ARG51, CYS2-ASN39 |
| HCGETSWQTGDFV | 5DRW | B | -85.9 | -1579.94 | SER6-GLN205, THR9-GLN205 |
|  | 1K1F | A | -29.14 | -366.08 | GLN8-ARG22 |
| KGTLEPEYF | 5DRW | B | -91.21 | -1767.94 | THR3-SER20, GLU7-SER10 |
|  | 1K1F | A | -35.76 | -629.66 | GLU5-GLN51 |
| KTVGELGDVRE | 5DRW | B | -37.47 | -1490.12 | THR2-PRO125, VAL3-SER127, GLU11-ASN144, GLY7-SER168, GLY7-SER182 |
|  | 1K1F | A | -54.36 | -686.76 | THR5-ARG25 |
| LTGTGVLTESNK | 5DRW | B | -40.46 | -1235.01 | THR4-TYR37, THR4-ASN39, SER10-GLN43 |
|  | 1K1F | A | -110.1 | -652.47 | GLU9-SER18, GLU9-SER41 |

For protein structure 5DRW, interaction and total energy of peptide and B-cell receptor were varied from −94.17 to −37.47 kcal/mol and −1656.88 to −1235.01 kcal/mol respectively. Whereas interaction and total energy for 1K1F were varied from −110.1 to −25 kcal/mol and −686.76 to −362.98 kcal/mol respectively.

## 4. Discussion

SARS-CoV-2 virus has infected more than four million people worldwide, and contagious nature of virus has imposed the biggest challenge of COVID-19 treatment and prevention. Therefore, vaccine design, development and production against COVID-19 diseases is an urgent requirement to protect people from the rising viral attacks. In practice, whole process of vaccine development takes several years to be completed [43]. But, integration of immunological understanding, high throughput genomics technologies, and bioinformatics tools and techniques can help us to design effective and safe vaccines in a short duration. In human research studies, vaccines were shown variable length of protection period such as chikungunya, rift valley fever virus, and measles, are about 30 years, 12 years, and 65 years, respectively [44]. In order to develop epitope-based vaccine, the surface glycoprotein is the primary focus because it is involved in the interaction between virus and human cell receptor, and contribute significant role in pathogenesis. But, other viral elements are also important to cause disease [45]. Information of expressed regions of viral genome is very important to identify potential vaccine candidates. In the present study, SARS-CoV-2 transcriptome was used for the molecular cataloguing of immunodominant epitopes, and result of performed analysis is summarized in Figure-5.

**Figure – 5.**
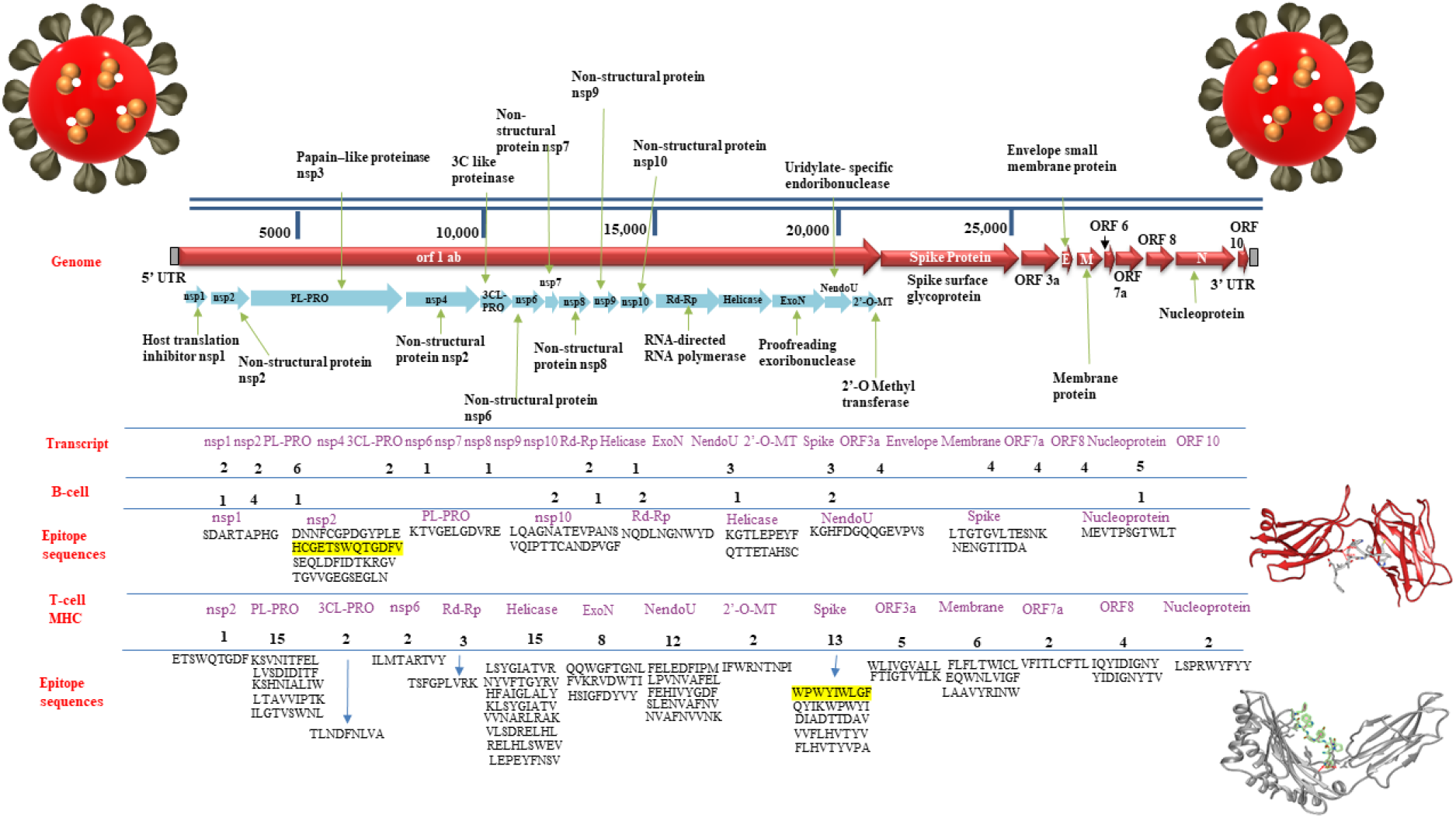
Overview of performed study: genome-wide transcriptome analysis of SARS-CoV-2, T- and B-cell epitopes, and docked complexes. Docked structures of B-cell receptor and MHC alleles were given at right-side and their T-and B-cell epitopes were highlighted in yellow colour.

In this study, we used RNA-seq data of SARS-CoV-2 infection in human cell lines NHBE and A549. We extracted the RNA-seq reads for SARS-CoV-2 genome and constructed a transcriptome to explore expressed regions of SARS-CoV-2 genome. In assembled transcripts, non-structural polyprotein 1ab related transcripts were found as most abundant transcripts, whereas nucleoprotein, membrane protein, SARS_X4 domain-containing protein, spike protein were also found in significant numbers. The largest region of SARS-CoV-2 genome (ORF1ab) was expressed various smaller proteins such as non-structural proteins (NSP1, NSP2, NSP3, NSP4, NSP6 and NSP8), helicase, uridylate-specific endoribonuclease, RNA-dependent RNA polymerase. Expression of polyprotein 1ab protein subcomponent (Table – 1) is clearly reflecting about disease initiation and progression such as expression of NSP1 protein was indicating the inhibition of host translation machinery by making NSP1-40S ribosome complex which can cause an endonucleolytic cleavage near the 5’ UTR of host mRNAs for degradation. NSP1 also facilitates viral gene expression in infected cells by suppressing host gene expression [46]. NSP2 is another expressed transcripts that may have a role in the alteration of host cell survival signalling pathway by interacting with host prohibitin (PHB) and prohibitin-2(PHB2)[47]. NSP3 is known as papain-like proteinase, which is responsible for N-terminus cleavage of the replicase polyprotein and involved in the assembly of virally-induced cytoplasmic doublemembrane vesicles together with NSP4, necessary for viral replication. NSP3 is an important viral molecular factor to suppress innate immune induction of type I interferon by blocking the phosphorylation, dimerization and subsequent nuclear translocation of host interferon regulatory transcription factor (IRF3), and also suppress NF-kappa-B signalling of host [48, 49]. NSP6 initiate induction of autophagosomes from host reticulum endoplasmic. But later, it reduces the expansion of phagosomes to stop delivery of viral components to lysosomes [50]. NSP8, together with NSP7 forms hexadecamer act as a primase to participate in a viral replication. Expression of helicase is required for the RNA and DNA duplex-unwinding activities [51].

Antigenicity and immunogenicity are the important parameters of epitopes selection. Antigenicity of epitopes represents the ability to bind or interact with B-cell or T-cell receptors. T-cell receptors recognize amino acid sequences of epitopes, when it binds with MHC molecules whereas in B-cell, B-cell receptor interact with these epitopes. To identify antigenic epitopes, all the epitopes were selected at a threshold antigenicity score greater than 1 (Table – 2). In our antigenicity analysis, epitopes KSVNITFEL (2.138), IQYIDIGNY (2.096), RELHLSWEV (2.260), and FTIGTVTLK (2.032) were containing high antigenicity. In comparison of antigenicity, immunogenic features of epitopes triggers the innate immune response, and later induces adaptive immune response. Antigenic epitopes may or may not have immunogenicity. But, all immunogenic epitopes will have antigenic potential [52]. In order to select best immunodominant epitopes, antigenicity (>=1) and immunogenicity (>=0.1), both criteria were used (Table-2) i.e. WPWYIWLGF (0.417), LSPRWYFYY (0.357), FLFLTWICL (0.354), FELEDFIPM (0.335), KSVNITFEL (0.330), IQYIDIGNY (0.304), QQWGFTGNL (0.281), LSYGIATVR (0.256), LVSDIDITF (0.254), and FVKRVDWTI (0.253). Epitopes with higher antigenicity and immunogenicity scores will have a higher probability of binding with T-and B-receptors to elicit an effective immune response. MHC proteins helps to distinguish cell own proteins from foreign proteins such as viruses and bacteria. Thus, the binding affinity of these epitopes to MHC protein is another very important criteria. MHC class I and II genes and alleles were predicted with lower IC_50_ values (IC_50_ =<200) to ensure a higher affinity of epitopes binding with MHC class I proteins. When MHC class I molecules binds to epitopes, immune system recognizes these epitopes as a foreign peptide, and the infected cell presents itself as an antigen-presenting cell for self-destruction. For better sensitivity, CD8^+^ T-cell epitopes should be generated from both structural and non-structural proteins because both types of proteins will be processed by infected cells in the cytoplasm, whereas structural proteins are of interest for CD4^+^ T-cell epitopes, as it might provide help to cognate interaction [53–55]. In this study, we selected CD4^+^ T-cell epitopes on the basis of CD8^+^ T-cell epitope core sequences to find out the best T-cell epitopes which might provide an immune response for both kinds of MHC classes. Twelve 15-mer MHC class II epitopes were selected which have core sequences of four 9-mer epitopes of MHC class I, and all four CD8^+^ cell epitopes belong to surface glycoprotein (Table – 3). Diverse repertoire of MHC molecules with the binding ability to a wide range of epitopes, and genetic variability among SARS-CoV clade antigens are the major scientific challenges to develop generalized vaccine. In order to address these two challenges, we performed conservation analysis of identified epitopes through IEDB resources and NCBI Blast, and epitope conservation were ensured among known sequences. IC_50_ threshold 500nM or lower values were suggested to selects strong binding affinity between MHC class protein and epitopes [56]. To ensure MHC class allele specificity, lower IC_50_ (=<200nM) values were considered for the selection of MHC alleles and epitopes. Genetic diversity of MHC molecules across the various ethnic groups worldwide is another the major limitation such as different MHC class alleles form different geographical region might be presented by a particular set of epitopes only. To understand demographic coverage of epitopes, population coverage analysis was performed through selected T-cell epitopes, and analysis revealed that approximately 89.44% and 93% average coverage can be achieved for world population and population of ethnic groups respectively (Figure-2, Supplementary file1: Table – S3).

In various B-cell research studies, particular antigen induces distinct class or subclass of antibodies such as schistosomiasis and filariasis induced a mixed response of IgE and IgG [57, 58]. In order to select distinct class of epitopes, sequence and structure-based dual approaches were used to identify B-cell epitope by using BepiPred-2.0 and ElliPro programs. 14 non-toxic, non cross-reactive, antigenic, and immunogenic B-cell epitopes were identified of different length (Table – 4). Predicted epitopes may or may not be a key feature of proteins because prediction methods, ignored epitope and receptor interaction, may be predicted only putative epitopes, which might lead to produce an antibody of no use. The real epitopes cannot be identified without considering the structural compatibility of complex formation [41]. Therefore, it became important to determine the structural coordinates of peptide-binding pockets to identify motifs for peptide binding. The specificities of different MHC class alleles and B-cell receptor’s phenotypes can be used to predict the recognition patterns of epitopes derived from antigens. The molecular docking approach was used to validate the interaction of three T-cell epitopes to most frequently occurring twenty-two MHC allele’s structures (Table – 5). Similarly, the top five B-cell epitopes were used to explore peptide interaction with two different kind of B-cell receptor proteins (5DRW and 1K1F). First protein structure, 5DRW was considered to evaluate binding affinity of peptides to BCR antibody light chain, whereas 1K1F was a Bcr-Abl oncoprotein and formed a tetramer through oligomerization. Monomer of 1K1F protein provided a basis to design an inhibitors to disrupt Bcr-Abl oligomerization[42]. Therefore, 1K1F was considered as control to compare the peptide binding affinity to B-cell receptor Fab region binding affinity (Table – 6). All peptides showed higher binding affinity to 5DRW than 1K1F except peptide KTVGELGDVRE (Supplementary file1: Table – S8). Docked complex of 1K1F and peptides was contained good interaction and total energy (−37.47, and −1490.12) and six hydrogen bonds (THR2-PRO125, VAL3-SER127, GLU11-ASN144, GLY7-SER168, GLY7-SER182). Hydrogen bond visualization of 1K1F protein and peptide were given in Figure-4.

**Figure – 4.**
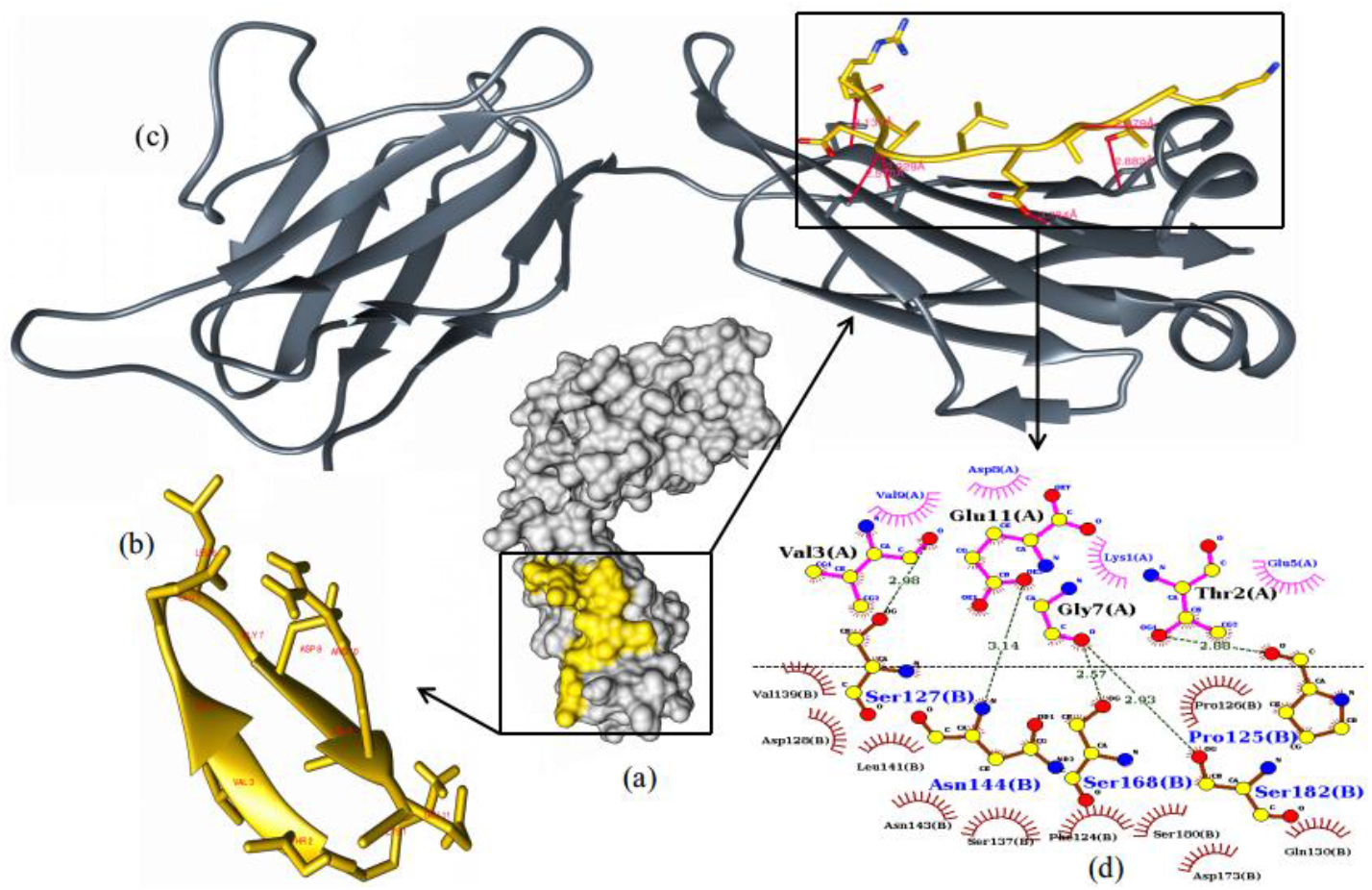
Molecular interaction of peptide (KTVGELGDVRE) and protein structure of Fab region of B-cell receptor (5DRW). a and c) cartoon structure of peptide and B-cell receptor. b) interaction of peptide and B-cell receptor (Hydrogen bonds in red colour). d) molecular level description of interaction through LIGPlot software (Hydrogen bonds in green colour)

The MHC genes and regions are one of the highly studied parts of human genome because it is highly associated with different diseases, immune responses and natural targets for molecular evolution, and very well characterized at functional levels [59]. Due to biomedical interest, MHC class gene expression profiling was performed through SARS-CoV-2 RNA-seq data, and HLA-A variants (HLA-A*01, HLA-A*02, HLA-A*03, HLA-A*11, HLA-A*24, HLA-A*25, HLA-A*26, HLA-A*30, HLA-A*31, HLA-A*32, HLA-A*33, HLA-A*34) were more expressed than HLA-B (HLA-B*08, HLA-B*15,HLA-B*18,HLA-B*37,HLA-B*44,HLA-B*45) and HLA-C variants (HLA-C*03, HLA-C*04, HLA-C*06, HLA-C*07, HLA-C*12, HLA-C*16). In MHC class II analysis, HLA-DMA*01, HLA-DMB*01, HLA-DOB*01, HLA-DPB1*08, HLA-DPB1*75, HLA-DQA1*01, HLA-DQB1*06, HLA-DRA*01 and HLA-DRB1*13 gene variants were highly expressed to present exogenous antigens to CD4^+^ T cells. Name of expressed MHC class genes and alleles, expression count (TPM) of alleles with respective to human cells were given in supplementary material file1 (Table – S8). Total, 92 gene variants of HLA-A*24 were expressed. Interestingly, very low or no expression for HLA-A*24 gene variants were observed for A549 cell line. In literature, prevalence of HLA-A*24 alleles was suggested as risk factors for severe H1N1 infection [60], and HLA-A*24:02 alleles were also reported to increase diabetes-associated risk together with HLA-B*39:01 gene [61]. A HLA-C allele, HLA-C*03:03, linked male infertility was also highly expressed in SARS-CoV-2 infected NHBE cell lines. The performed study on semen quality was reported that presence of HLA-C*03:03 allele was increased two fold in human papillomavirus virus infected individuals [62]. Three genetic variant of HLA-B*08:01 genes, myasthenia gravis autoimmune disease characterized by muscle weakness and abnormal fatigability were also highly expressed in NHBE cell lines [63]. In present, a combination of malaria and AIDS drugs are in use for the treatment of COVID-19. So, it would be interesting to explore malaria and HIV associated MHC class allele’s in SARS-CoV-2 transcriptome. Therefore, all the MHC class genes were analyzed and filtered, and a list of expressed MHC class alleles for malaria and HIV were generated by using IEDB resources along with expression counts for human cell lines (Supplementary file1: Table – S9). HLA-A*02:01:131, HLA-A*02:01:160 and HLA-A*03:01:01:02N were expressed for malaria and HIV in NHBE cell lines. SARS related two HLA genes (HLA-A*23:01:03, HLA-A*23:01:31) were expressed, and 34 HLA-A*24:02 were expressed for HIV gag polyprotein in NHBE cell lines. In our analysis, HLA-A*23:01 HLA-A*24:02 and HLA-A*02:01 were predicted for proposed T-cell epitopes, and molecular interaction studies were also performed between T-cell epitopes and revealed MHC alleles.

## 5. Conclusion

This study has high scientific relevance to understand immune responses of SARS-CoV-2 in the scarcity of experimental resources. Performed study has provided the extremely useful information about the expressed region of SARS-CoV-2 genome, potential T- and B-cell epitopes, molecular interaction of identified epitopes to receptors through various bioinformatics approaches, and gene expression of MHC class I and II genes. Expressed regions of SARS-CoV-2 genome and putative expressed targets of human immune response will facilitate vaccine related research studies. Proposed epitopes are possessing T- and B-cell selectivity, nontoxicity, higher population coverage, and significant interaction with MHC class I and II genes, and B-cell receptors. However, the presented list of T-and B-cell epitopes is an outcome of computational analysis. But, all the epitopes were identified from transcriptome data of SARS-CoV-2 infection in human cell-lines. Therefore, these epitopes have high potential to reflect SARS-CoV-2 immune response, and become vaccine candidates after experimental validation.

## Author’s contributions

SKK: conceived the study, planned the data processing and analysis, and written first draft of manuscript.

VB and SS: performed the docking studies.

SC and SG: contributed in biological data interpretation, manuscript writing, and generation of figures and tables.

Final manuscript is read and approved by all the authors.

## Acknowledgment

We are grateful to the Director NIAB for providing support to carry out the study. Bioinformatics facility, NIAB is acknowledged for the bioinformatics data analysis. We would also thankful to Aakash Chawade and Pallavi Chauhan for useful discussion and suggestions.

## Supplementary information

Supplementary file 1(Table S1-S9): Meta data for used transcriptome, detail description of T- and B-cell epitopes, detail description docking studies and gene expression analysis of MHC-alleles

Supplementary file 2: Verification of genuine hydrogen bonds involved in the interaction through cavity prediction.

## Data availability

All the used RNA-seq data is available at NCBI SRA under the project accession number PRJNA615032.

