## Supplementary_file2 for "SARS-CoV-2 transcriptome analysis and molecular cataloguing of immunodominant epitopes for multi-epitope based vaccine design"

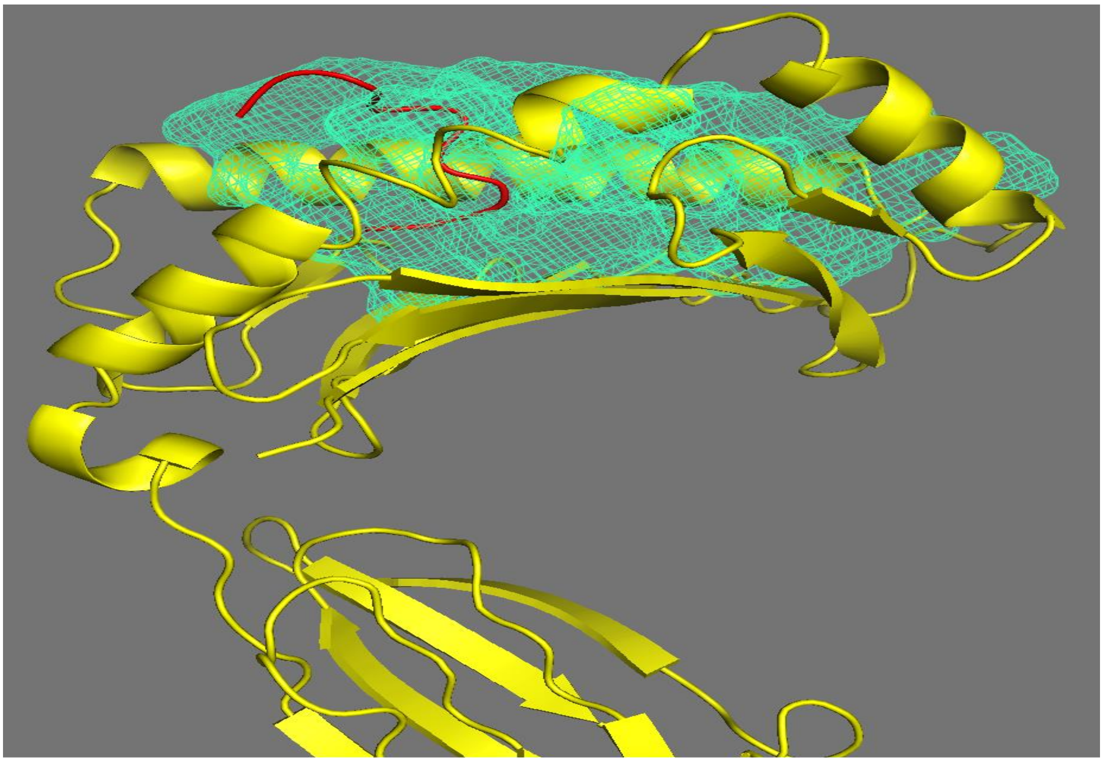

Figure -1. Hydrogen bond verification through MHC (HLA-B*57:01) and peptide (WPWYIWLGF) docked complex. Hydrogen bonds and interacting residues were residing in-side cavity (TRP6-ASN66, PHE9-TYR59, PHE7-TRY99, TRP6-PHE9, and TRP1-GLU63) and all the hydrogen bonds were covered in cavity number 2.

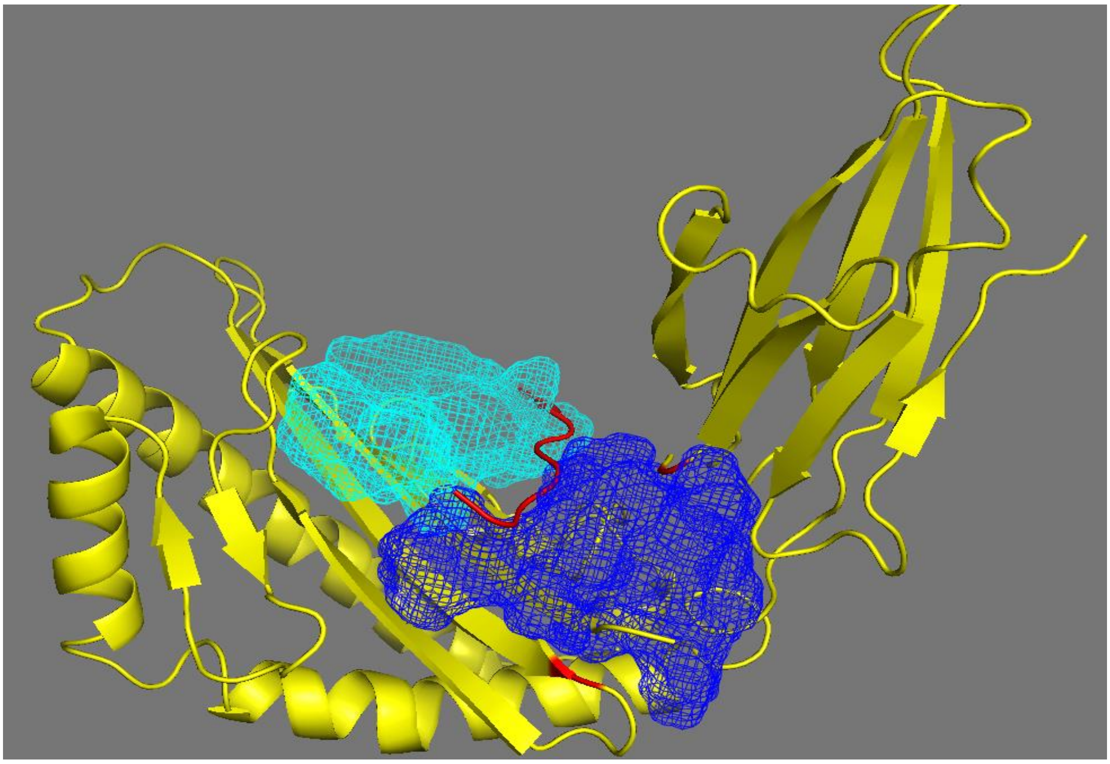

Figure –2. Hydrogen bond verification through MHC (HLA-B*35:01) and peptide (WPWYIWLGF) docked complex. Hydrogen bonds and interacting residues were residing in-side cavity (TYR4-APG6, TRP6-TYR27, LEU7-TYR27, TYR4-THR233, TRP1-HIS113, and TYR4-THR233) and all the hydrogen bonds were covered in cavity number 2 and 3.

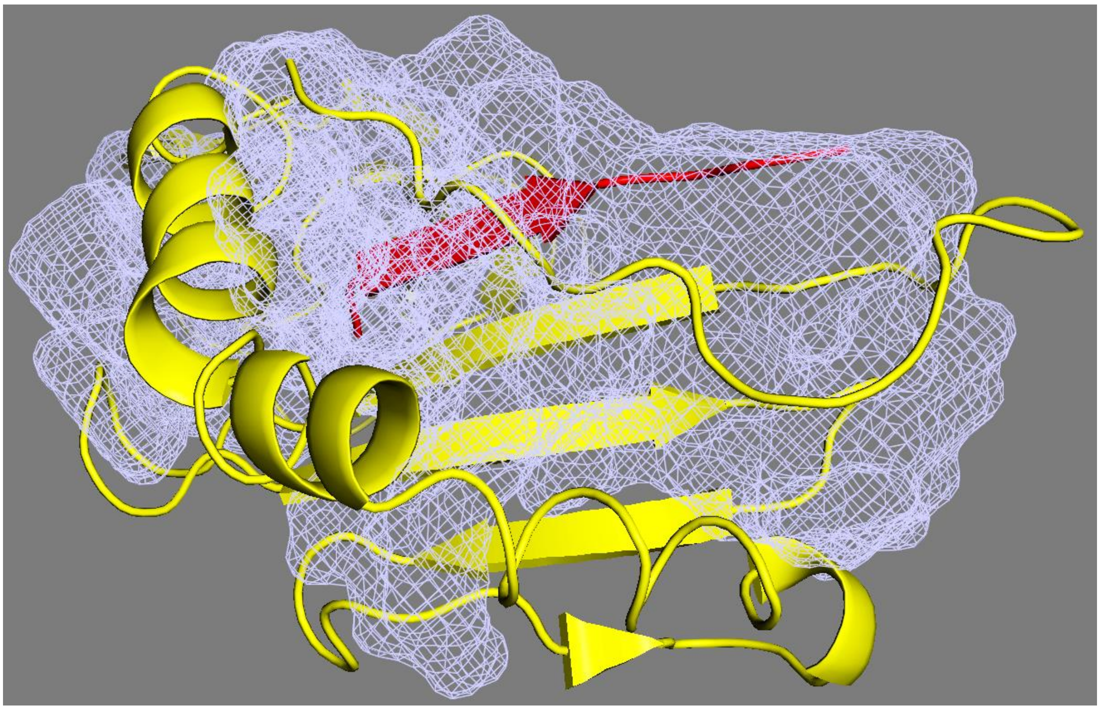

Figure -3. Hydrogen bond verification through MHC (HLA-A*23:01) and peptide (FLHVTYVPA) docked complex. Hydrogen bonds and interacting residues were residing in-side cavity (LEU2-CYS64, VAL4-PHE262, TYR6-LEU260), and all the hydrogen bonds cover in cavity number 1.

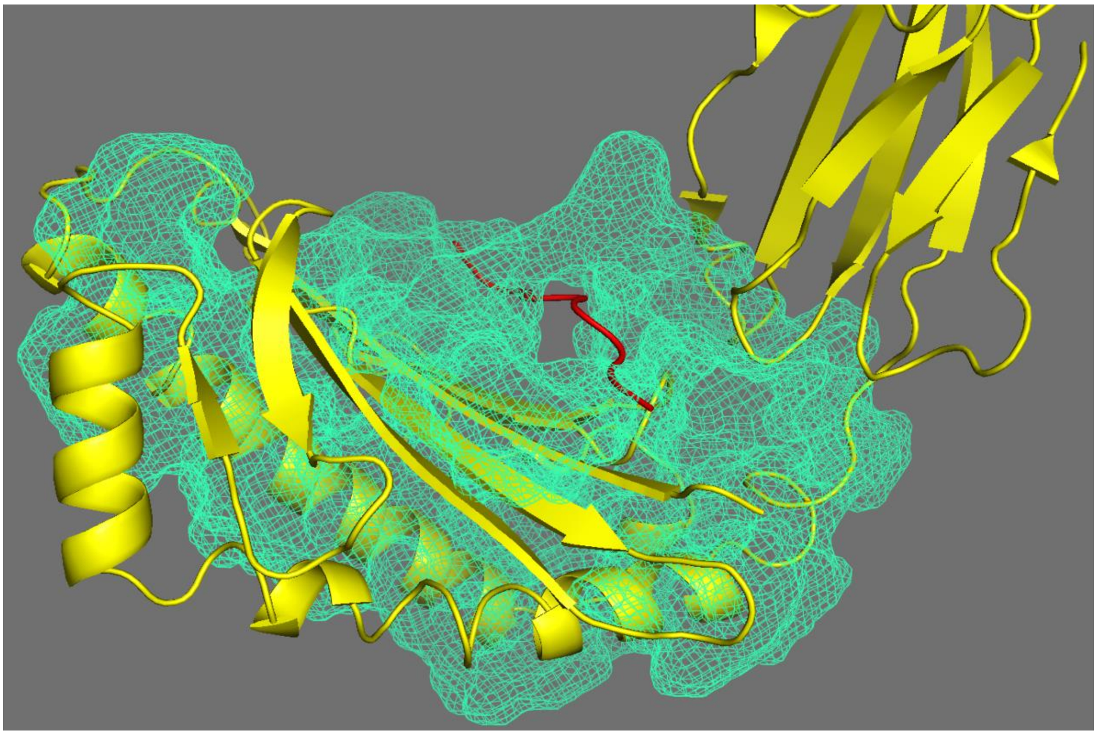

Figure -4. Hydrogen bond verification through MHC (HLA-A*11:01) and peptide (VVFLHVTYV) docked complex. Hydrogen bonds and interacting residues were residing in-side cavity (LEU4-TYR27, LEU4-ARG6, TYR8-SER4, SER4-VAL9), and all the hydrogen bonds cover in cavity number 1.

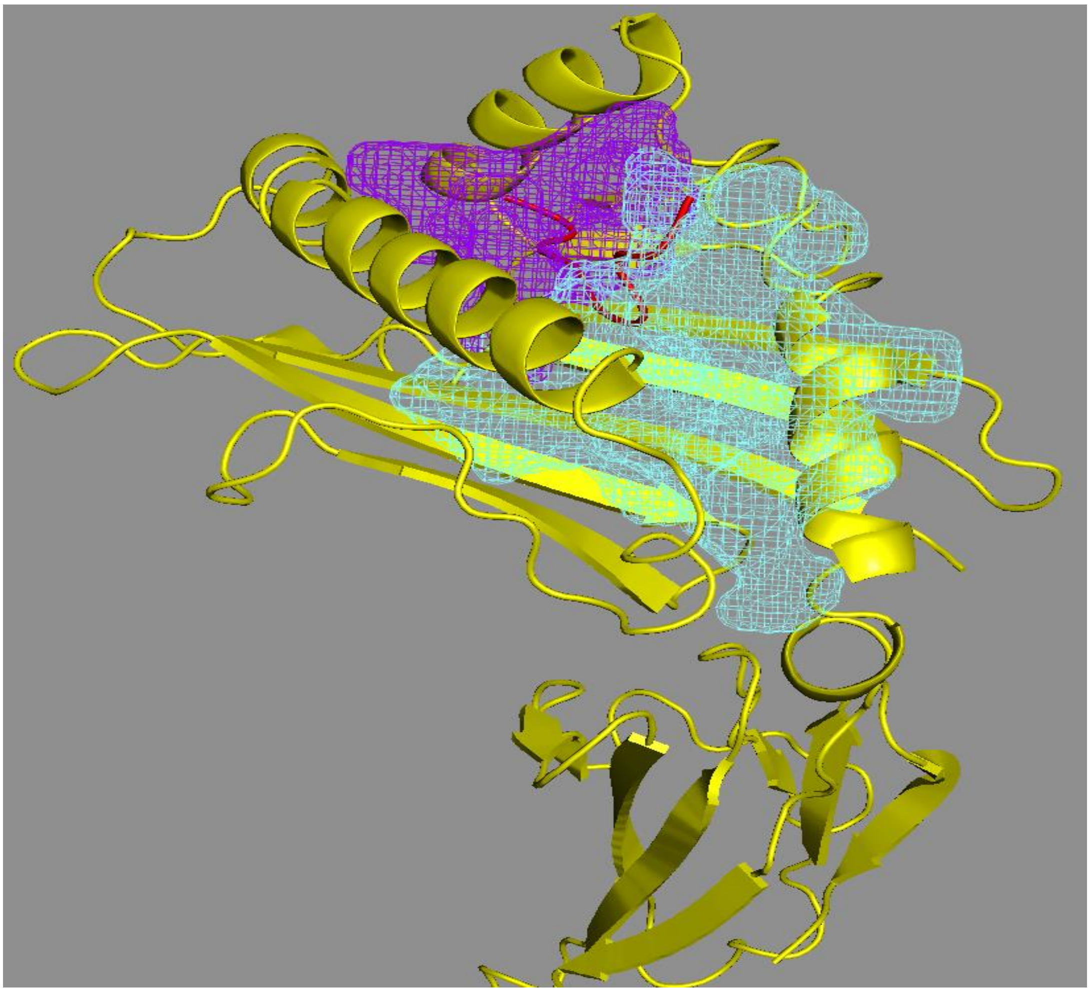

Figure -5. Hydrogen bond verification through MHC (HLA-A*11:01) and peptide (WPWYIWLGF) docked complex. Hydrogen bonds and interacting residues were residing in-side cavity (TYR4-CYS164, LEU7GLN156, PHE9-ASP116, GLY8-ASP116) and all the hydrogen bonds cover in cavity number 2 and 6.
